# Sequential dynein effectors regulate axonal autophagosome motility in a maturation-dependent pathway

**DOI:** 10.1101/2020.11.01.363333

**Authors:** Sydney E. Cason, Peter Carman, Claire Van Duyne, Juliet Goldsmith, Roberto Dominguez, Erika L.F. Holzbaur

## Abstract

Autophagy is a degradative pathway required to maintain neuronal homeostasis. Neuronal autophagosomes form constitutively at the axon terminal and mature via lysosomal fusion during dynein-mediated transport to the soma. How the dynein-autophagosome interaction is regulated during maturation is unknown. Here, we identify a series of handoffs between dynein effectors as autophagosomes transit along the axons of primary neurons. In the distal axon, JIP1 initiates autophagosomal transport, while autophagosomes in the mid-axon require HAP1 and Huntingtin for motility. We demonstrate that HAP1 is a bonafide dynein activator, binding the dynein-dynactin complex via canonical and noncanonical interactions. JIP3 is found on most axonal autophagosomes but specifically regulates the transport of acidified autolysosomes. Inhibiting autophagosomal transport disrupts maturation, while inhibiting autophagosomal maturation perturbs the association and function of dynein effectors. Thus maturation and transport are tightly linked. These results describe a novel maturation-based dynein effector handoff on neuronal autophagosomes that is key to autophagosomal motility, cargo degradation, and the maintenance of axonal health.

**Summary:** Neuronal autophagosomes form in the distal axon and mature via fusion with lysosomes during their dynein-driven transport to the soma. Dynein is regulated on autophagosomes by distinct effector proteins—JIP1, HAP1, and JIP3—depending on location and autophagosomal maturity. In this sequential pathway, transport and maturation state are tightly linked to maintain neuronal health.

## Introduction

Neurons are some of the longest living cells in the body, differentiating during development and persisting throughout the lifetime of the organism. To maintain homeostasis and cellular function, neurons require continuous clearance and recycling of aged proteins and organelles. Macroautophagy (hereafter, autophagy) is a major conserved degradation pathway essential for neuronal health (Kulkarni et al., 2018). Genetic disruption of the autophagy pathway in neurons is sufficient to induce neurodegeneration in mice (Hara et al., 2006; Komatsu et al., 2006). Deficits in neuronal autophagy are observed in most common neurodegenerative diseases including Alzheimer’s disease, and disease-causing mutations occur in proteins implicated in the autophagy pathway, including PINK1 and Parkin in Parkinson’s disease, optineurin and TBK1 in ALS, and Huntingtin in Huntington’s disease (Wong and Holzbaur, 2015).

Neurons are the longest cells in the body reaching up to 1 m in humans and are highly polarized; these unique features pose challenges for intracellular trafficking pathways including neuronal autophagy. Neuronal autophagosomes form de novo at presynaptic sites and axon terminals (Maday et al., 2012; Stavoe et al., 2016; Neisch et al., 2017). Once formed, autophagosomes mature by fusing with late endosomes and lysosomes during their transport to the soma along axonal microtubules (Maday et al., 2012). To traverse the distance from the axon terminal to the soma, where degradatively active lysosomes are enriched, axonal autophagosomes undergo highly processive retrograde motility. However, the regulatory mechanisms responsible for this transport are not well understood.

Microtubules (MTs) are organized in a polarized array in axons, with minus-ends directed toward the soma (Heidemann et al., 1981). Dynein is the primary minus-end directed microtubule motor in neurons, transporting nearly all retrograde-moving cargoes. Effector proteins link dynein to specific cargo and induce the formation of an activated motile complex made up of dynein and its obligate partner complex dynactin (Gill et al., 1991; McKenney et al., 2014; Zhang et al., 2017; Reck-Peterson et al., 2018; Olenick and Holzbaur, 2019). A new class of dynein effectors known as activating adaptors (1) stabilize the interaction between dynein and dynactin, (2) help align dynein’s two motor domains into an active parallel conformation required for efficient stepping along the microtubule, (3) link dynein-dynactin to specific cargos, and (4) can bring together multiple dynein dimers to increase force and processivity (Urnavicius et al., 2015, 2018; Grotjahn et al., 2018; Elshenawy et al., 2019). Activating adaptors that regulate the motility of some cargos, e.g. Rab6-positive vesicles and signaling endosomes (Matanis et al., 2002; Olenick et al., 2019), have been identified; however, the specific cargo adaptor and/or activator required for highly processive autophagosomal motility has not yet been determined.

The maturation process alters autophagosomal membrane composition through fusion with lysosomes, further complicating the regulation of dynein motors bound to this cargo. We investigated candidate proteins known to associate with autophagosomes and/or lysosomes. c-Jun N-terminal kinase (Jnk)-interacting protein 1 (JIP1) interacts directly with the autophagosomal membrane protein MT-associated protein 1 light chain 3 (LC3), dynactin, and the plus-end-directed MT motor kinesin-1 (Fu and Holzbaur, 2013; Fu et al., 2014). JIP1 is required for the initiation of the long-range transport of autophagosomes from the distal tip of neurites of dorsal root ganglion (DRG) neurons (Fu et al., 2014). Huntingtin (Htt) and its interacting partner Htt-associated protein 1 (HAP1) have also been implicated in autophagosomal transport in DRG neurons, though the underlying mechanism is unknown (Wong and Holzbaur, 2014). Htt interacts directly with dynein, while HAP1 binds both dynactin and kinesin-1 (Caviston et al., 2007; Engelender et al., 1997; Li et al., 1998; McGuire et al., 2006). Jnk-interacting protein 3 (JIP3; *Drosophila* Sunday driver/SYD; *C. elegans* UNC-16) has mostly been studied in the context of endolysosomal transport, but was recently shown to facilitate the dynein-mediated transport of autophagosomes along the axon in *C. elegans* (Marchesin et al., 2015; Hill et al., 2019). Structurally unrelated to JIP1, JIP3 interacts with kinesin-1, dynein, and dynactin (Cavalli et al., 2005; Arimoto et al., 2011; Cockburn et al., 2018; Vilela et al., 2019), suggesting that JIP3 may also contribute to the regulation of autophagosomal motility in mammalian neurons.

Here, we examine the trafficking of autophagosomes along the axons of primary rat hippocampal neurons. We identify multiple dynein effectors bound to autophagosomes during transit along the axon. Surprisingly, the functional requirement for each of these effectors changes with subaxonal location and with the maturation state of the the autophagosome. JIP1 colocalizes primarily with nascent autophagosomes in the distal axon, consistent with its role in initiation of processive retrograde transport. HAP1 robustly associates with autophagosomes in the mid-axon and is required for autophagosomal motility specifically within that region. We used complementary in vitro and cellular assays to demonstrate that HAP1 functions as an activating adaptor of dynein and identified both canonical and noncanonical determinants that regulate this interaction. Finally, we find that JIP3 colocalizes with autophagosomes throughout the axon, but primarily regulates the motility of more mature autolysosomes in the proximal axon. While inhibition of motility has been shown to disrupt autophagosomal maturation (Wong and Holzbaur, 2014), we find genetic or pharmacological inhibition of autophagosomal maturation perturbs interactions between autophagosomes and dynein effectors, indicating that motility and maturation state are tightly linked. Together, these results illustrate a novel coordination mechanism between autophagosomal maturation state and motor regulation by effector proteins.

## Results

### Multiple dynein effectors associate with the autophagosomal membrane in axons

To investigate the association of dynein effectors with axonal autophagosomes, we assayed the colocalization of JIP1, HAP1, and JIP3 with the autophagosome marker LC3B in the axons of live and fixed primary rat hippocampal neurons. First, we co-expressed Halo-tagged versions of each effector with mCherry(mCh)-EGFP-LC3B and imaged neurons at 6–8 days in vitro (DIV) using live-cell confocal microscopy. All three effectors localized to the axon as puncta that colocalized and comigrated with LC3-positive (LC3+) puncta (Fig. 1, A-C). When we co-expressed EGFP-LC3 with all three candidates—BFP-JIP1, mCh-HAP1, and Halo-JIP3—we frequently observed simultaneous colocalization of all candidates with LC3+ puncta (Fig. S1 A).

**Figure 1.**
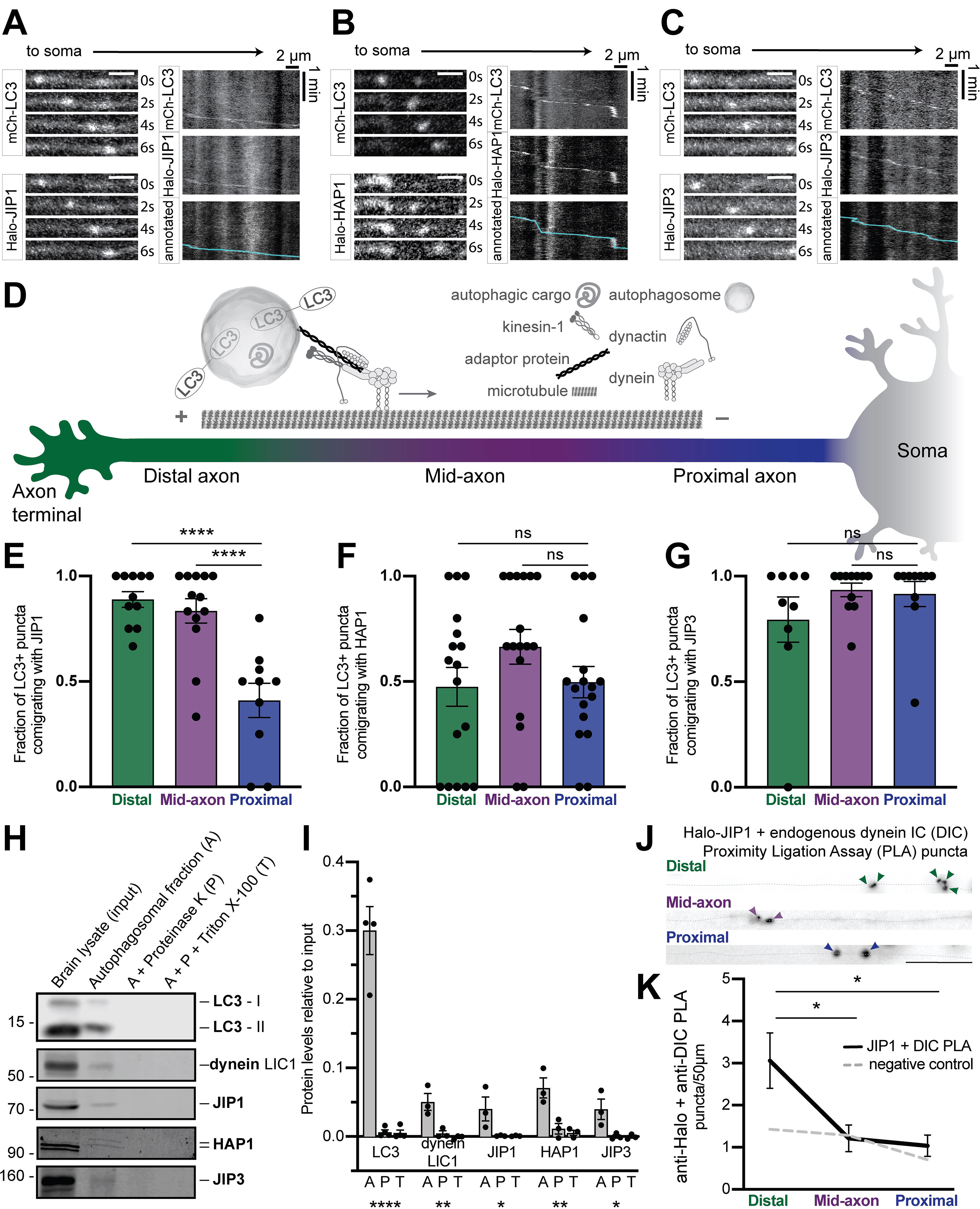
Multiple dynein effectors associate with the autophagosomal membrane in axons. (A-C) Time series and kymographs from separate LC3+ autophagosomes demonstrating comigration with JIP1 **(A)**, HAP1 **(B)**, and JIP3 **(C)**. **(D)** Schematic illustrating neuronal subregions and axonal transport of autophagosomes. (E-G) Quantification of LC3+ puncta comigrating with JIP1 **(E)**, HAP1 **(F)**, JIP3 **(G)** in different subaxonal regions. *n* = 9-17 neurons; one-way ANOVA (*p* < 0.0001) with Tukey’s multiple comparisons test (Distal v. mid-axon and Mid-axon v. proximal, *p* < 0.0001). **(H)** Immunoblotting of autophagosomal isolation illustrating enrichment of LC3, dynein, and candidate effectors on the autophagosomal surface. **(I)** Quantification of enrichment in the autophagosomal fraction, displayed as relative to brain lysate (input). *n* = 3-4 preps; one-way ANOVA for each protein; LC3 both isoforms (*p* < 0.0001), dynein light intermediate chain 1 (LIC1; *p* = 0.0056), JIP1 (*p* = 0.0497), HAP1 both isoforms (*p* = 0.0058), JIP3 (*p* = 0.0277). **(J)** Representative micrograph showing PLA puncta for endogenous dynein intermediate chain (DIC) with Halo-JIP1 along different subregions of axon (dotted gray line). Arrows indicate proximity ligation assay (PLA) puncta. Scale bar, 10 µm. **(K)** Quantification of PLA puncta per 50 µm axon. Dashed gray line, negative control (missing primary antibody). *n* = 7 neurons; one-way ANOVA (*p* = 0.0085) with Tukey’s multiple comparisons test (Distal v. mid-axon, *p* = 0.0236; Mid-axon v. proximal, *p* = 0.0131). Bars throughout figure show mean ± SEM. ns, not significant; *, *p* < 0.05; **, *p* < 0.01; ****, *p* < 0.0001.

Previous work on JIP1 demonstrated that it specifically promoted autophagosomal motility in the distal tips of neurites (Fu et al., 2014), suggesting dynein may require distinct effectors to drive cargo motility in different subaxonal regions. To test this hypothesis, we investigated the colocalization of these effectors with LC3B-positive (LC3+) puncta in the distal (<100 µm from the axon tip), proximal (<100 µm from the soma), and mid-axon (Fig. 1 D; Fu et al., 2014; Olenick et al., 2019). JIP1 colocalized to a significantly higher degree with LC3+ puncta in the distal axon and the mid-axon, and showed less colocalization in the proximal axon (Fig. 1 E). HAP1 and JIP3, in contrast, colocalized to a similar extent with LC3+ puncta in all regions (Fig. 1, F and G).

To further assess the association of these effectors with LC3+ autophagosomes, we performed proximity ligation assays (PLA), used to detect proteins within 40 nm of one another (Alam, 2018). Neurons expressing low levels of Halo-JIP1, Halo-HAP1, or Halo-JIP3 were fixed and stained with primary antibodies to Halo-tag and to endogenous LC3B, then secondary antibodies conjugated to complementary oligonucleotides were added to assess their physical proximity (Fig. S1 B). All three effectors were closely apposed to LC3 in the axon, as measured by density of fluorescent oligonucleotide puncta (Fig. S1, C-H). However, none demonstrated subaxonal specificity.

We then isolated autophagosomes from mouse brain via sequential ultracentrifugation, adding Gly-Phe-β-naphthylamide to inactivate and deplete lysosomal vesicles and thus enhance the integrity of autophagosome-associated proteins (Strømhaug et al., 1998; Maday et al., 2012). To differentiate between autophagosomal cargo engulfed within the organelle and endogenous membrane-associated proteins associated with the cytosolic face of the organelle, we treated the autophagosome-enriched fraction with either Proteinase K alone to degrade exposed proteins or with Proteinase K and Triton X-100 to degrade all proteins. The autophagosomal fraction was appropriately enriched for the lipidated form of LC3, LC3-II, and depleted of markers for other organelles (Fig. S1, I-K). Both dynein and JIP1 were significantly enriched on the cytosolic face of isolated autophagosomes (Fig. 1, H-I). This enrichment is consistent with previous findings that JIP1 binds directly to LC3 via an LC3-interacting region (LIR) motif (Fu et al., 2014). JIP3 was also enriched on the autophagosome surface (Fig. 1, H-I). JIP3 may interact with the autophagosome via its binding partner Arf6, which has been shown to be important for autophagosomal trafficking in zebrafish (Montagnac et al., 2009; George et al., 2016). We also found both HAP1 (Fig. 1, H-I) and its interacting partner Htt (Fig. S1, I-K) associated with the cytosolic surface of the autophagosome. This is consistent with previous data showing HAP1 and Htt enriched in an autophagosome-enriched fraction prepared from mouse brain (Wong and Holzbaur, 2014). While HAP1 is not known to interact directly with any autophagosomal membrane proteins, Htt has been shown to coimmunoprecipitate with LC3B from HEK293T cell lysate and contains a validated binding site (aa3037-3040) for another member of the LC3 family, GABARAPL1 (Ochaba et al., 2014). This suggests that Htt may link HAP1 to the autophagosomal membrane.

Finally, we asked whether the candidate proteins were closely apposed to dynein in the axon. PLA between the Halo-tagged effectors and endogenous dynein (dynein intermediate chain [DIC]) revealed that JIP1–dynein complexes were significantly enriched in the distal axon where the number of puncta was roughly 3-fold higher than in other regions (Fig. 1, J-K). Consistent with the subaxonal specificity seen here, JIP1 is required for initiating autophagosomal motility in the distal axon (Fu et al., 2014). HAP1 and JIP3 exhibited proximity to dynein throughout the axon (Fig. S1, L-O) and, unlike JIP1, did not show regional specificity. These results suggest that multiple adaptors— JIP1, HAP1, and JIP3—interact with dynein on axonal autophagosomes.

### HAP1 regulates autophagosomal motility specifically in the mid-axon

To test the function of HAP1 in the motility of axonal autophagosomes, we used siRNA to knockdown (KD) HAP1 in neurons transduced with LC3B-GFP and imaged live at 6-8 DIV. HAP1 was depleted ∼85% in PC12 cells using this siRNA (Fig. S2, A-B). As expected, the majority of LC3+ puncta demonstrated robust retrograde motility towards the soma under control conditions (Fig. 2 A). We investigated the effect of HAP1 KD in the distal (Fig. 2B), mid-(Fig. 2, A and C), and proximal (Fig. 2 D) axonal regions to test subaxonal specificity. HAP1 depletion induced a strong phenotype in the mid-axon where the majority of LC3+ puncta exhibited stationary or bidirectional motility, defined as events moving <10 µm during the video (Fig. 2, A and C). A subset of autophagosomes continued to move in the retrograde direction, which may be due to incomplete KD or scaffolding by other dynein effectors. To confirm specificity of KD, we expressed siRNA-resistant wild- type Halo-HAP1 (HAP1^WT^) in KD neurons, which rescued autophagosomal motility to control levels (Fig. 2, E-F). Interestingly, the inhibitive effect of HAP1 KD on autophagosomal motility was limited to the mid-axon; HAP1 KD had no effect on autophagosomal motility in the distal (Fig. 2 B) or proximal (Fig. 2 D) regions.

**Figure 2.**
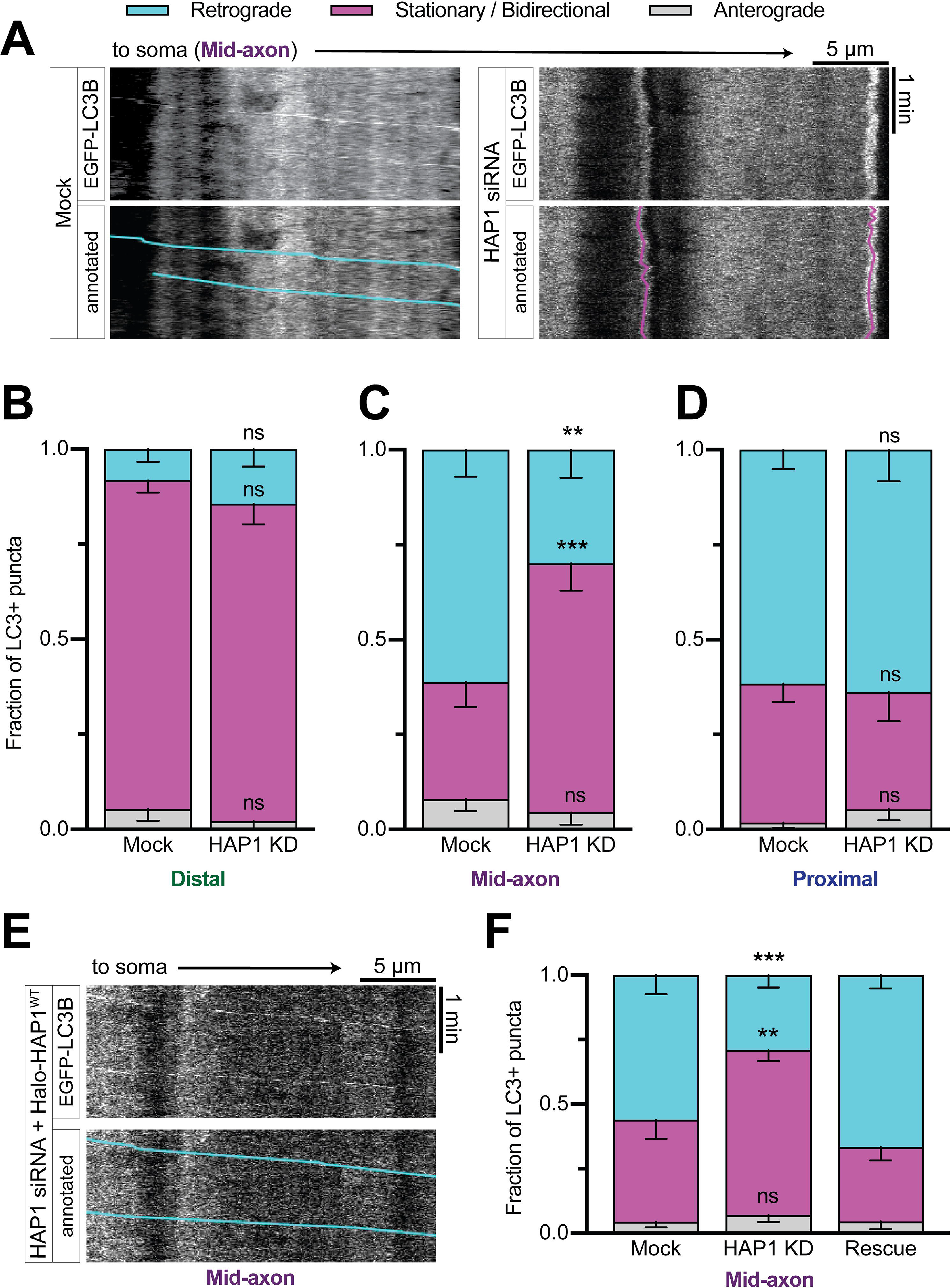
HAP1 regulates autophagosomal motility specifically in the mid-axon. **(A)** Representative kymographs from the mid-axon of a mock-transfected (control) neuron and a neuron transfected with HAP1 siRNA. Kymographs depict distance on the x-axis and time on the y-axis. Annotated kymographs mirror the above kymographs with the LC3+ puncta paths pseudo-colored for easier visualization. (B-D) Quantification of LC3+ puncta motile behavior in the distal **(B)**, mid-**(C)**, and proximal **(D)** axonal subregions. *n* = 8-21 neurons. Two-way ANOVA for each with Sidak’s multiple comparisons test (Mid-axon Retrograde Mock v. HAP1 KD, *p* = 0.0033; Mid-axon Stationary/Bidirectional Mock v. HAP1 KD, *p* = 0.0009). (E-F) Representative kymograph **(E)** and quantification **(F)** from the mid-axon of a neuron transfected with HAP1 siRNA and siRNA-resistant Halo-HAP1^WT^. Two-way ANOVA with Tukey’s multiple comparisons test (Retrograde Mock v. HAP1 KD, *p* = 0.0005; Stationary/Bidirectional Mock v. HAP1 KD, *p* = 0.0020). Bars throughout figure show mean ± SEM. Symbols (ns, not significant; **, *p* < 0.01; ***, *p* < 0.001) indicate Sidak’s or Tukey’s test comparing to Mock.

Knockdown of Htt (Fig. S2, A-B) likewise increased the stationary/bidirectional fraction in the mid-axon with no effect on motility in the distal or proximal axon (Fig. S2, C-F). Expression of siRNA-resistant mCh-Htt rescued autophagosomal motility (Fig. S2, G-H). Together our data show that the HAP1-Htt complex is required for autophagosomal motility exclusively in the mid-axon of hippocampal neurons.

### HAP1 binds dynein-dynactin via dynein activating adaptor motifs

We next queried the mechanism by which HAP1 regulates autophagosomal motility. Based on structural predictions, we hypothesized HAP1 directly activates the dynein-dynactin complex. Cryo-EM studies of known dynein activators in complex with dynein and dynactin have identified a conserved length of coiled-coil necessary to dock along dynactin’s filament of actin related protein 1 (Arp1) and occupy the dynein-dynactin interface (Hodgkinson et al., 2005; Urnavicius et al., 2015, 2018). Sequence analysis of HAP1 using 4 different in silico coiled-coil prediction programs (Lupas et al., 1991; Trigg et al., 2011; Vincent et al., 2013; Ludwiczak et al., 2019) identified an extended coiled-coil segment (Fig. 3 A). Assuming a 3.5-residue periodicity, the predicted coiled-coil domain is likely to extend 33 nm (Truebestein and Leonard, 2016), the appropriate length to dock along the Arp1 filament (35 nm).

**Figure 3.**
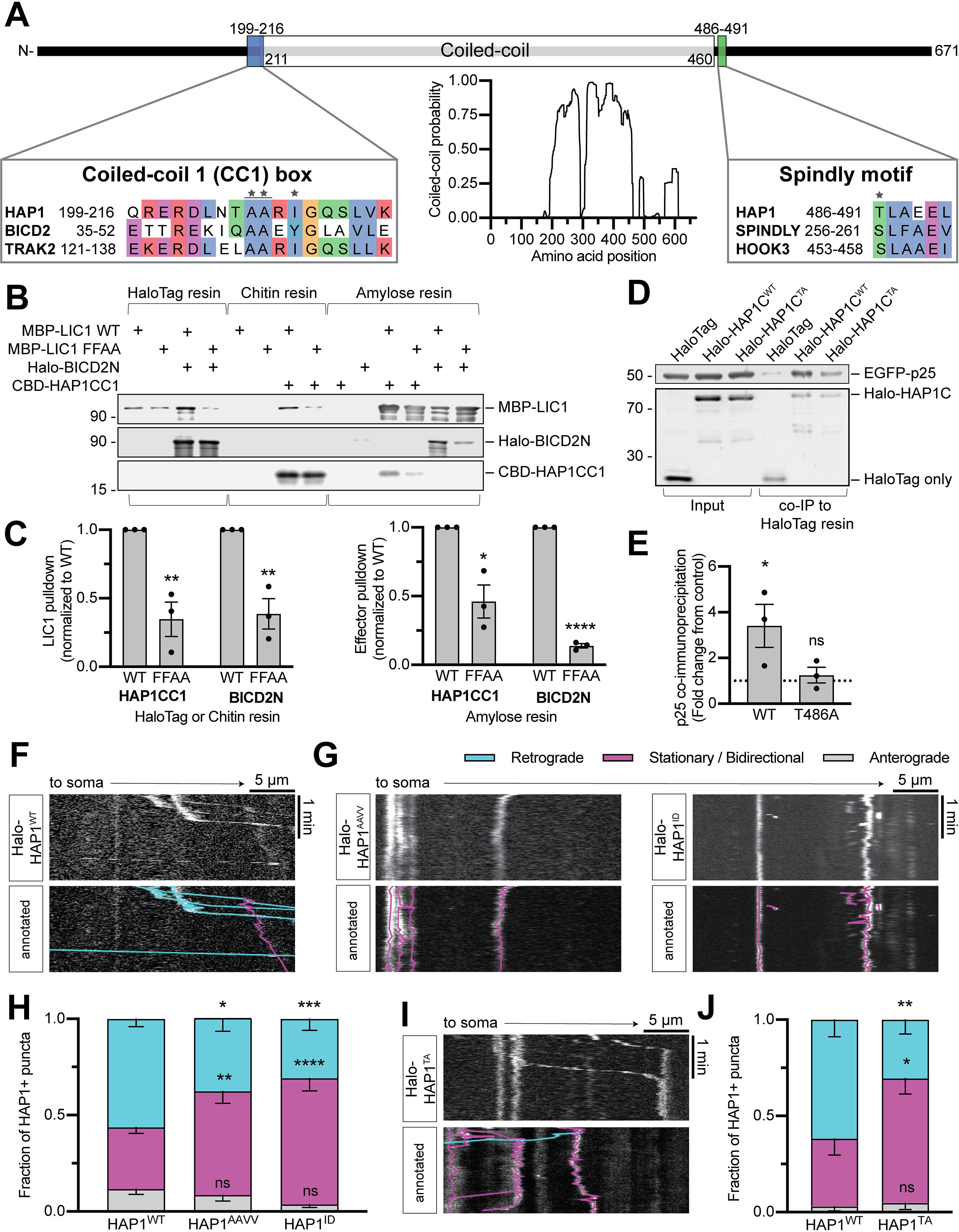
HAP1 binds dynein-dynactin via dynein activating adaptor motifs. **(A)** Schematic illustrating the domain architecture of HAP1. Coiled coil probability was calculated using predictions from 4 different programs. Amino acids in the sequence alignments are colored using the clustal color scheme where conserved. Stars indicate point mutants for (D-J). **(B)** Immunoblot showing purified HAP1CC1 (aa 168-261) or BICD2N pulls down purified dynein LIC1^WT^ but not mutant LIC1^FFAA^ and vice versa. **(C)** Quantification of purified protein pulldowns. Pulldowns were normalized to the LIC1^WT^ condition and tested via unpaired t test (two-tailed; LIC1 pulldown by HAP1CC1 WT v. FFAA, *p* = 0.0066; LIC1 pulldown by BICD2N WT v. FFAA, *p* = 0.0051; HAP1CC1 pulldown by LIC1 WT v. FFAA, *p* = 0.0109; BICD2N pulldown by LIC1 WT v. FFAA, *p* < 0.0001). *n* = 3 pulldown experiments from *n* = 2 independent protein purifications. (D-E) Example immunoblot **(D)** and quantification **(E)** showing HAP1C^WT^ (aa 470-671) construct co-expressed in Cos7 cells with EGFP-p25 co-immunoprecipitates p25, while HAP1C^TA^ does not. *n* = 3 repeats; results were normalized to p25 pulled down by HaloTag only (negative control; dotted black line); one-way ANOVA of the 3 conditions (*p* = 0.0488) with Dunnett’s multiple comparisons test to compare both HAP1C conditions to the control condition (HAP1C^WT^, *p* = 0.0456; HAP1C^TA^, *p* = 0.9351). (F-J) Example kymographs and quantification of HAP1+ puncta motile behavior in the mid-axon demonstrating HAP1^WT^ **(F)** moves primarily retrograde while HAP1 CC1 box **(G-H)** and HAP1 spindly **(I-J)** mutants exhibit stationary behavior. *n* = 11-16 neurons; two-way ANOVA with Bonferroni’s multiple comparisons test (Retrograde WT v. AAVV, *p* = 0.0333; Stationary/Bidirectional WT v. AAVV, *p* = 0.0099; Retrograde WT v. ID, *p* = 0.0009; Stationary/Bidirectional WT v. ID, *p* < 0.0001; Retrograde WT v. TA, *p* = 0.0063; Stationary/Bidirectional WT v. TA, *p* = 0.0109). Bars throughout figure show mean ± SEM. ns, not significant; *, *p* < 0.05; **, *p* < 0.01; ***, *p* < 0.001; ****, *p* < 0.0001.

The coiled-coil region in HAP1 is flanked by two conserved sequence motifs found in other dynein activating adaptors (Fig. 3 A). The motif preceding the coiled-coil domain has been labeled the coiled-coil 1 (CC1) box (Gama et al., 2017). The CC1 box directly interacts with the first alpha helical region of dynein subunit light intermediate chain 1 (LIC1; Lee et al., 2018; Celestino et al., 2019; Lee et al., 2020). To test whether HAP1’s predicted CC1 box motif has a similar function, we performed pull-down experiments using purified recombinant HAP1 spanning amino acid (aa) residues 168-261 (HAP1CC1) and purified recombinant LIC1. The N-terminus of known dynein activator Bicaudal-D 2 (BICD2N) was used as a positive control. Both HAP1CC1 and BICD2N pulled down LIC1 (Fig. 3, B-C). Parallel experiments using an LIC1 mutant with two phenylalanine-to-alanine substitutions known to disrupt LIC1-activator binding (Lee et al., 2018) markedly decreased pulldown (Fig. 3, B-C). In the reverse experiment, wild type but not mutant LIC1 pulled down HAP1CC1 and BICD2N (Fig. 3, B-C). HAP1 thus binds dynein’s LIC1 using the same motif as BICD2 and other CC1-box-containing activators.

The other conserved motif following the extended coiled-coil region was originally described in the activating adaptor Spindly and is thus termed the “Spindly motif” (Gama et al., 2017). The Spindly motif mediates the interaction between dynein activators and the pointed-end complex of dynactin’s Arp1 filament (Hodgkinson et al., 2005; Gama et al., 2017). To test this motif, we engineered a threonine-to-alanine HAP1 point mutant (T486A) predicted to deleteriously impact dynactin binding based on work in Spindly (Fig. 3 A; Gassmann et al., 2010; Gama et al., 2017). To hone in specifically on this latter motif, we generated a HAP1 C-terminal construct (aa 470-671) with (HAP1C^TA^) and without (HAP1C^WT^) the mutation. We co-transfected Cos7 cells with the dynactin pointed-end complex protein EGFP-p25 and either HaloTag alone, Halo-HAP1C^WT^, or Halo-HAP1C^TA^, then immunoprecipitated the lysate using HaloTag beads. HAP1C^WT^ co-immunoprecipitated (co-IP) significantly more p25 than the negative control HaloTag only, but HAP1C^TA^ did not (Fig. 3, D-E). Therefore, HAP1 contains a bonafide Spindly motif for binding the dynactin pointed-end complex.

Finally, to test the function of these binding motifs in cells, we overexpressed Halo-HAP1 in primary neurons and acquired videos along the axon, where the microtubules are polarized with plus-ends out. Overexpressed Halo-HAP1 formed HAP1-positive (HAP1+) puncta in the axon that moved primarily in the retrograde direction, indicating dynein activity (Fig. 3 F). To test the CC1 box motif, we introduced two substitutions (A207V, A208V) to alter conserved alanine residues known to be important for LIC1 binding in other proteins (HAP1^AAVV^; Fig. 3 A; Schlager et al., 2014; Gama et al., 2017; Lee et al., 2020). We also generated a second CC1 box point mutant based on the recent BICD2-LIC1 structure (Lee et al., 2020): we introduced a single isoleucine-to-aspartic acid mutation (I210D) where a hydrophobic residue is essential to create a hydrophobic groove for accomodating LIC1 Helix-1 (HAP1^ID^; Fig. 3 A). We overexpressed Halo-HAP1^ID^, HAP1^AAVV^ (Fig. 3, G-H), or the full-length Spindly motif mutant HAP1^TA^ (Fig. 3, I-J) into primary neurons and found all three mutants demonstrated significantly less retrograde motility compared to HAP1^WT^ (Fig, 3, H and J). Hence, both the dynein-and dynactin-binding regions in HAP1 are required for dynein motilty in cells.

### HAP1 contains a novel conserved p150^Glued^ binding site

Apart from the interaction between HAP1 and the dynactin pointed-end complex, a previous study identified a direct interaction between dynactin subunit p150^Glued^ and HAP1 via a yeast two-hybrid screen (Engelender et al., 1997). To test this interaction, we co-transfected Cos7 cells with FLAG-p150^Glued^ and either HaloTag only, Halo-HAP1^WT^, Halo-HAP1^TA^, or BICD2N-Halo then immunoprecipitated the lysate. Both HAP1^WT^ and HAP1^TA^ were capable of co-immunoprecipitating p150^Glued^, indicating p150^Glued^ binding and pointed-end complex binding are independent (Fig. 4, A-B). Conversely, BICD2N was incapable of co-immunoprecipitating p150^Glued^ above negative control (HaloTag only) levels. In the reverse pulldown, p150^Glued^ co-immmunoprecipitated both HAP1 constructs but not BICD2N or HaloTag alone (Fig. 4, A-B).

**Figure 4.**
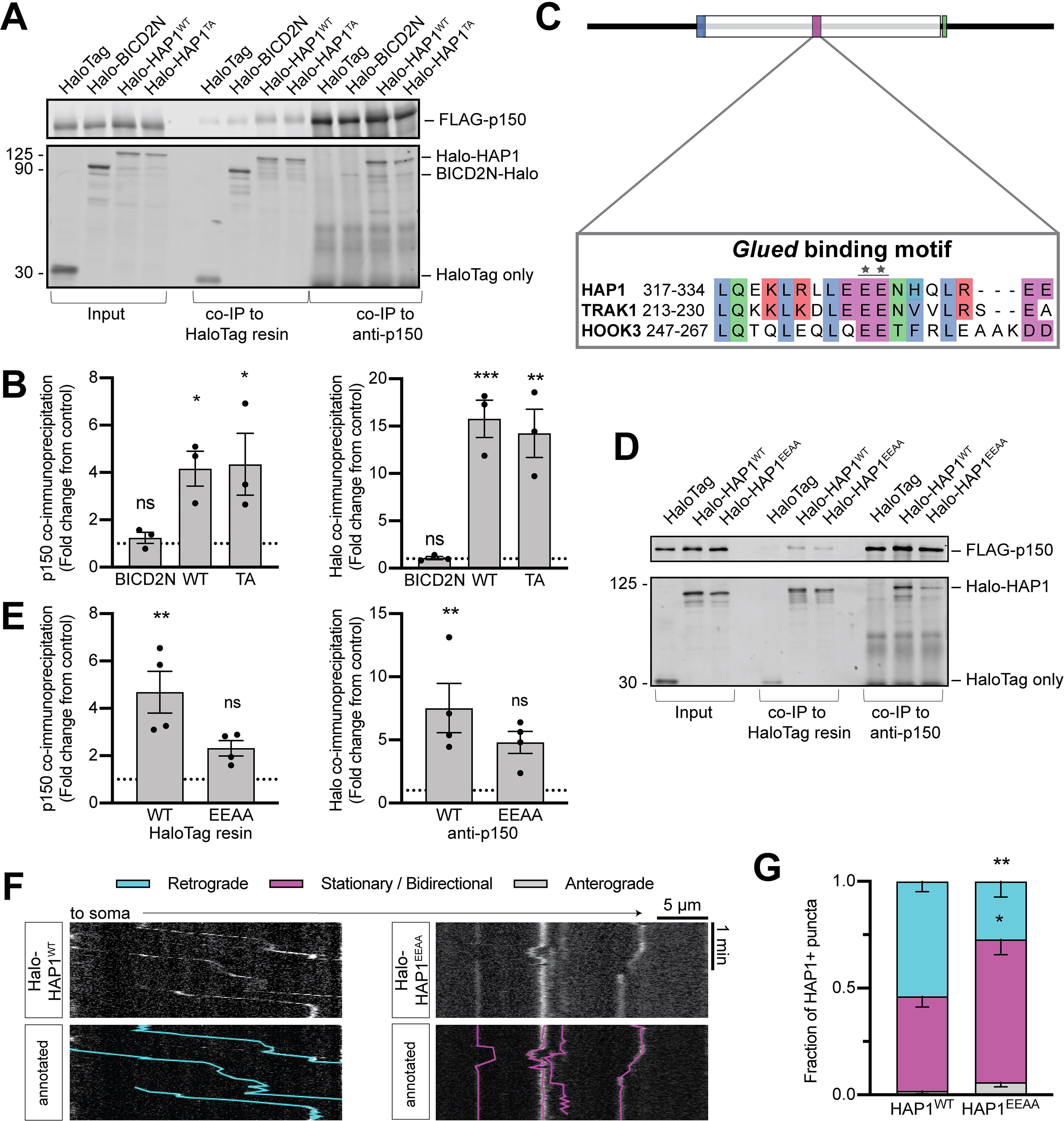
HAP1 contains a novel conserved dynactin p150^Glued^ binding site. (A-B) Example immunoblot **(A)** and quantification **(B)** showing both HAP1 constructs, but not BICD2N, bind dynactin subunit p150^Glued^ (p150) and coimmunoprecipitate together from Cos7 cells. *n* = 3 repeats; results were normalized to HaloTag only (negative control; dotted black line); one-way ANOVA of the 3 conditions (left, *p* = 0.0218; right, *p* = 0.0002) with Dunnett’s multiple comparisons test to compare all to the control condition (Left: HAP1^WT^, *p* = 0.0452; HAP1^TA^, *p* = 0.0351. Right: HAP1^WT^, *p* = 0.0005; HAP1^TA^, *p* = 0.0011). **(C)** Schematic showing sequence alignment for the *Glued* binding motif. Stars indicate point mutant. (D-E) Immunoblotting **(D)** and quantification **(E)** reveals HAP1^WT^ but not the *Glued* motif mutant HAP1^EEAA^ binds p150. *n* = 4 repeats; results were normalized to HaloTag only (negative control; dotted black line); one-way ANOVA of the 3 conditions (left, *p* = 0.0030; right, *p* = 0.0144) with Dunnett’s multiple comparisons test to compare to the control condition (Left: HAP1^WT^, *p* = 0.0018; HAP1^EEAA^, *p* = 0.2036. Right: HAP1^WT^, *p* = 0.0085; HAP1^EEAA^, *p* = 0.0996). (F-G) Example kymographs **(F)** and quantification **(G)** of HAP1+ puncta motile behavior in the mid-axon demonstrating HAP1^WT^ moves primarily retrograde while HAP1^EEAA^ exhibits more stationary behavior. *n* = 13-15 neurons; two-way ANOVA with Bonferroni’s multiple comparisons test (Retrograde WT v. EEAA, *p* = 0.0019; Stationary/Bidirectional WT v. EEAA, *p* = 0.0103). Bars throughout figure show mean ± SEM. ns, not significant; *, *p* < 0.05; **, *p* < 0.01; ***, *p* < 0.001.

Conservation analysis indicates that the HAP1 region spanning aa residues 280-445, initially identified in the yeast two-hybrid screen, is well conserved across species (Fig. S2 I). This region is also ∼30% conserved within the evolutionarily-related TRAK family of proteins (Fig. S2 J; Lumsden et al., 2016) and ∼25% conserved within the HOOK family of known dynein activating adaptors, which have been shown to coimmunoprecipitate strongly with p150^Glued^ (Fig. S2 J; Olenick et al., 2016). The region is less conserved in other dynein activators, including Spindly and the BICD family of proteins (Fig. S2 K). We identified a highly conserved motif within this region and engineered a double glutamate-to-alanine HAP1 mutant construct to test p150^Glued^ binding (E326-327A; HAP1^EEAA^; Fig. 4 C). We co-transfected Cos7 cells with FLAG-p150^Glued^ and either HaloTag only, Halo-HAP1^WT^, or Halo-HAP1^EEAA^ then immunoprecipitated the lysate. As before, HAP1^WT^ co-immunoprecipitated p150^Glued^ and vice versa, but HAP1^EEAA^ did not (Fig. 4, D-E). Thus, we identify a novel *Glued* binding motif within HAP1 that mediates an interaction with dynactin subunit p150^Glued^. To test the function of this motif in cells, we transfected our *Glued* motif HAP1 mutant into primary neurons to observe its motile behavior (Fig. 4 F). The mutant displayed significantly less retrograde motility than HAP1^WT^, indicating the importance of this binding site for the formation of a motile dynein complex (Fig. 4 G). We therefore propose that both the HAP/TRAK family and the HOOK family possess an essential dynactin binding site that is independent from pointed-end-complex binding and not present in other well-characterized dynein activators.

### HAP1 activates dynein motility

Based on our biochemical data, we hypothesized that HAP1 functions as an activating adaptor for dynein. To test this hypothesis, we assayed the ability of HAP1 to redistribute organelles in live COS7 cells using an induced dimerization assay. Our cell-permeant dimerizer is made up of trimethoprim (TMP) conjugated to a HaloTag ligand and induces dimerization between two proteins wherein one is tagged with dihydrofolate reductase (DHFR) and the other contains a HaloTag (Ballister et al., 2015). We used peroxisomes as a model organelle because they are generally immotile and uniformly distributed under control conditions, making them ideal to observe changes in motility (Smith and Aitchison, 2013). We co-expressed PEX3-GFP-Halo and either HAP1-mCh-DHFR or, as a positive control, BICD2N-mCh-DHFR then assessed the redistribution of peroxisomes when the dimerizer was added (Olenick et al., 2016). Recruitment of either HAP1 (Fig. 5 A) or BICD2N (Fig. S3 A) to peroxisomes induced their retrograde motility and eventual clustering in the perinuclear region, near the MT organizing center. Perinuclear clustering was not seen in vehicle-treated cells (Fig. S3 B). HAP1 thus induces minus-end-directed organelle motility in a cellular assay.

**Figure 5.**
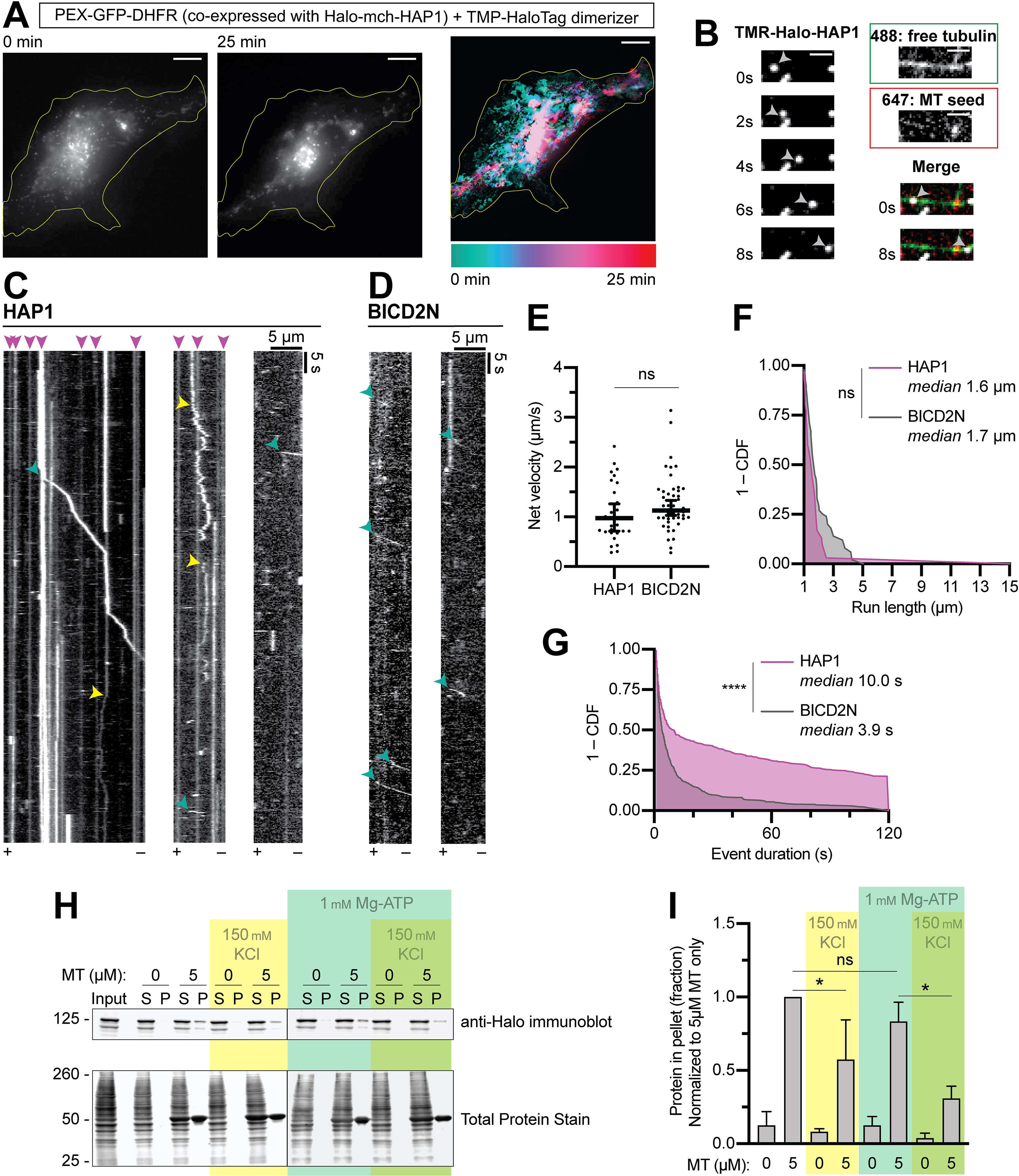
HAP1 activates dynein motility and microtubule binding. **(A)** Peroxisome recruitment assay wherein peroxisomes (PEX-GFP-DHFR) were tethered to Halo-mCh-HAP1 via a dimerizer and imaged over the following 25 min to show relocalization to the cell center. Left, initial (0 min) and concluding (25 min) stills of peroxisomes. Right, max time projection pseudocolored by frame (color scale below). Yellow line indicates cell outline. Scale bar, 10 µm. **(B)** Example micrographs from single molecule motility TIRF assay showing TMR-labeled Halo-HAP1 traversing a microtubule (MT) from the dynamic plus-end to the minus-end, indicated by the 647-labeled MT seed. Single stills from both tubulin channels are shown (488, 647) as well as the first and last frames merged (all three channels). Scale bar, 2 µm. (C-D) Example kymographs showing dynein-directed motility of HAP1 **(C)** and BICD2N **(D)** puncta in the lysate-based motility assay. Kymographs all represent full microtubules with the plus-end (+) and minus-end (-) labeled at the bottom. Arrows represent motile minus-end-directed events (teal), diffusive events (yellow), and sustained MT binding events (pink). (E-G) Quantifications of single molecule motility assay showing the average velocity **(E)**, run length **(F)**, and stationary event duration **(G).** *n* = 30-51 runs (E-F) or 457-471 events (G) in 3 independent trials on 57 total MT each. Welch’s two-tailed t test (E, *p* = 0.1969) or Mann-Whitney test (F, *p* = 0.0568; G, *p* < 0.0001). Y-axes in (F-G) represent 1 - the cumulative distribution function (CDF); X-axis in (F) begins at 1 µm because only runs ≥ 1µm were included in our analyses; X-axis in (G) ends at 120 s, the length of videos acquired. (H-I) Blotting **(H)** and quantification **(I)** of cell extract MT pelleting assay. *n* = 3 independent assays; one-way ANOVA (*p* < 0.0001) followed by Tukey’s multiple comparisons test. HAP1 is shown to bind MT in a largely ATP-independent fashion (*p* = 0.6691). Increasing ionic strength (150 mM KCl) decreased pellet fraction significantly (without ATP, *p* = 0.0075; with ATP, *p* = 0.0011) but not to 0 µM MT levels. Bars throughout figure show mean ± SEM. ns, not significant; *, *p* < 0.05; ****, *p* < 0.0001.

To more directly test the ability of HAP1 to activate dynein, we performed lysate-based in vitro single molecule motility assays using total internal reflection fluorescence (TIRF) microscopy (Ayloo et al., 2014). Cos7 cell lysates containing TMR-labelled HAP1 or BICD2N were introduced into flow chambers with MTs. HAP1 interacts with both kinesin and dynein so motility direction was key (McGuire et al., 2006; Twelvetrees et al., 2010). To determine MT polarity, we bound MT seeds (labeled with HiLyte 647 tubulin) to the coverslip then flowed in GTP and free tubulin (labeled with differently colored HiLyte 488 tubulin) and imaged MT dynamics; plus ends were identified by their higher rates of both growth and catastrophe. Both HAP1 (Fig. 5, B-C) and our positive control BICD2N (Fig. 5 D) induced minus-end-directed runs (teal arrows) with similar average velocities (1.1 µms^-1^) and run lengths (1.6 µm) to previous reports of dynein motility (Fig. 5, E-F; Olenick et al., 2016; Lee et al., 2020).

BICD2N exhibited almost exclusively (90%) minus-end-directed runs, while HAP1 displayed about 30% plus-end-directed runs, not surprising given the interaction between HAP1 and kinesin. However in both primary axons and in vitro, HAP1 demonstrates majority minus-end-directed motility. We assessed HAP1 activation of kinesin by performing a kinesin MT recruitment assay in TIRF with full-length kinesin-1 (Fig. S3, C-D). Although HAP1 has been shown to activate kinesin in coordination with another effector, GRIP1 (Twelvetrees et al., 2019), we found HAP1 could not activate kinesin independently or with Htt in this assay.

In addition to processive runs, HAP1 complexes displayed diffusive events (yellow arrows; Fig. 5 C) and frequent long immotile MT binding events, often lasting the entire length of the video (pink arrows; Fig. 5, C and G). These sustained MT binding events and diffusive runs, not seen in the BICD2N condition, are similar to those observed recently with purified TRAK1 in vitro (Henrichs et al., 2020). TRAK1, which is related to HAP1, was found to directly bind microtubules (Henrichs et al., 2020), leading us to hypothesize that the frequent non-processive events observed in our motility assay were due to HAP1 MT binding. While we were unable to purify soluble HAP1 to test direct MT binding, we performed a cell-extract-based MT pelleting assay in which Cos7 lysate expressing Halo-HAP1 was incubated with 5 µM MT under varying buffer conditions. HAP1 robustly pelleted with MT, and this pelleting was unaffected by addition of Mg-ATP (1 mM; Fig. 5, H-I) which caused the dynein-dynactin complex to disassociate from the MT (Fig. S3, E-F). High ionic strength buffer (150 mM KCl) decreased but did not eliminate HAP1 pelleting, suggesting the interaction is partially dependent on electrostatic interactions. TRAK1 microtubule binding provides an additional anchor to the microtubule during kinesin motility, allowing the motor complex to navigate crowded microtubule surfaces and increase run length (Henrichs et al., 2020). HAP1 may likewise use its microtubule interaction to prevent the motile complex from disassociating from the microtubule, which would stall autophagosomal transport during the long trip through the axon. Together, these observations indicate HAP1 can bind MTs and activate dynein, but not kinesin, motility.

### HAP1 drives autophagosomal transport by activating dynein

We next querried whether the ability of HAP1 to activate dynein motility contributes to autophagosomal transport in axons. We co-transfected our Halo-HAP1 constructs with EGFP-LC3 into primary neurons and acquired 3 min videos in the mid-axon. LC3+ puncta in neurons expressing HAP1^WT^ demonstrated primarily retrograde motile behavior (Fig. 6 A), while overexpression of either CC1 box mutant HAP1^AAVV^ or HAP1^ID^ (Fig. 6, B-C) or the *Glued* motif mutant HAP1^EEAA^ (Fig. 6, E-D) induced a dominant negative effect, significantly reducing the number of retrograde-moving LC3+ puncta. Similar to the results from the HAP1 KD, expression of the mutants had a dominant negative effect on LC3+ puncta motility in the mid-axon but not the proximal axon (Fig. S3, G-K).

**Figure 6.**
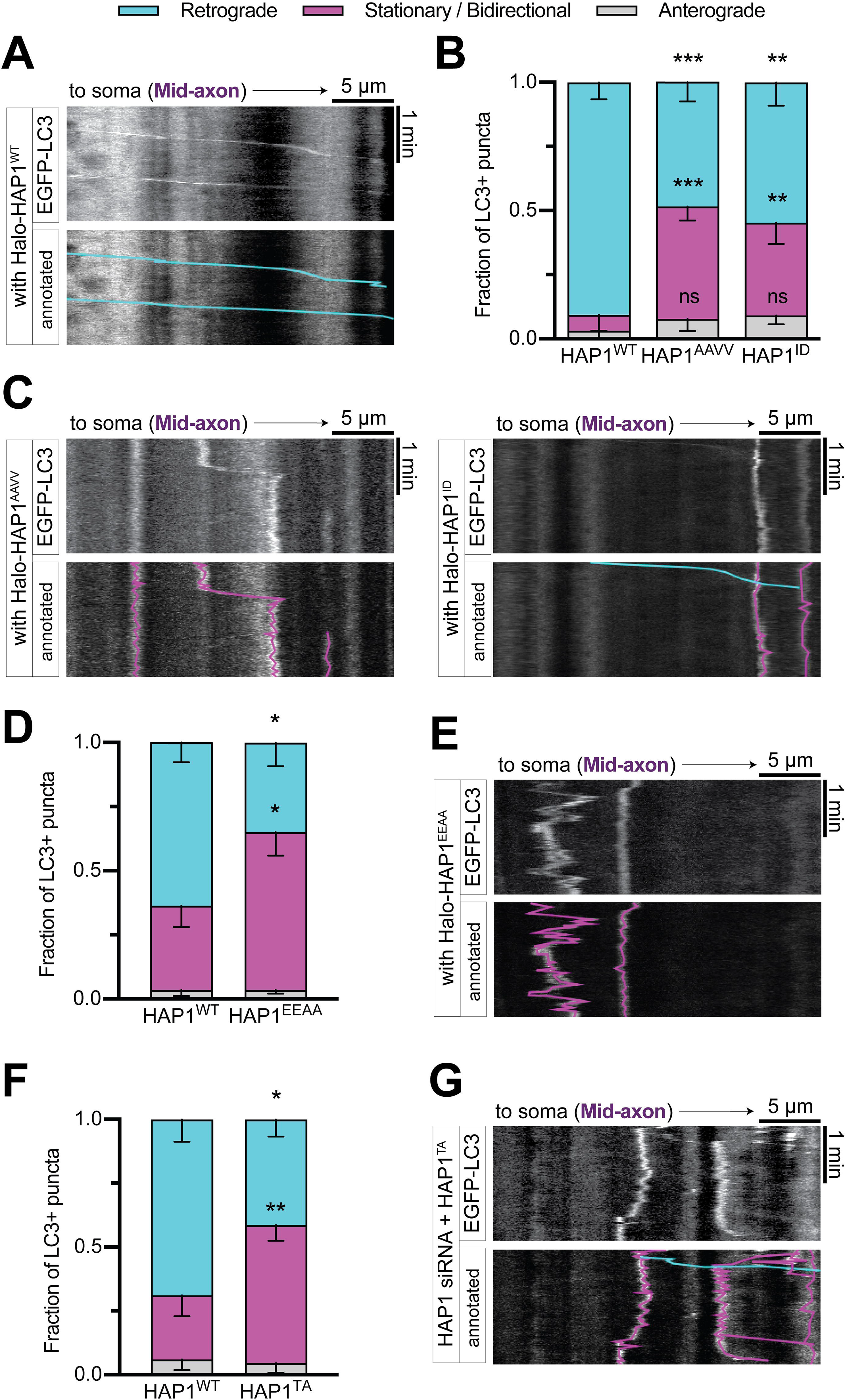
HAP1 drives autophagosomal transport by binding to dynein and dynactin. **(A)** Example kymograph illustrates the typical motility of EGFP-LC3+ puncta in the presence of overexpressed Halo-HAP1^WT^. (B-C) Quantification **(B)** and example kymographs **(C)** showing dominant negative effect of HAP1 CC1 box mutants (HAP1^AAVV^, HAP1^ID^) on LC3+ puncta motility. *n* = 8-12 neurons; two-way ANOVA with Bonferroni’s multiple comparisons test (Retrograde WT v. AAVV, *p* = 0.0002; Stationary/Bidirectional WT v. AAVV, *p* = 0.0009; Retrograde WT v. ID, *p* = 0.0013; Stationary/Bidirectional WT v. ID, *p* = 0.0091). (D-E) Quantification **(D)** and example kymograph **(E)** showing dominant negative effect of HAP1 *Glued* motif mutant (HAP1^EEAA^) on LC3+ puncta motility. *n* = 14-15 neurons; two-way ANOVA with Bonferroni’s multiple comparisons test (Retrograde WT v. EEAA, *p* < 0.0001; Stationary/Bidirectional WT v. EEAA, *p* = 0.0002). (F-G) Quantification **(F)** and example kymograph **(G)** showing nonmotile behavior of LC3+ puncta in cells transfected with both HAP1 siRNA and the HAP spindly mutant (HAP1^TA^). *n* = 16 neurons; two-way ANOVA with Bonferroni’s multiple comparisons test (Retrograde WT v. TA, *p* = 0.0071; Stationary/Bidirectional WT v. TA, *p* = 0.0075). Bars throughout figure show mean ± SEM. ns, not significant; *, *p* < 0.05; **, *p* < 0.01; ***, *p* < 0.001. Scale bars, 5 µm. All data from the mid-axon.

The Spindly mutant HAP1^TA^ had a less marked negative effect on LC3+ motility, which we hypothesized might be due to interaction with endogenous HAP1 (Fig. S3, L-M). To examine the requirement of the Spindly motif for autophagosome motility, we expressed siRNA-resistant Halo-HAP1^TA^ in neurons treated with HAP1 siRNA. In the absence of endogenous HAP1, Halo-HAP1^WT^ was able to induce retrograde motility of LC3+ puncta, but Halo-HAP1^TA^ failed to do so (Fig. 6, F-G). LC3+ puncta in neurons transfected with both HAP1 siRNA and HAP1^TA^ demonstrated primarily nonmotile behavior in the mid-axon (Fig. 6, F-G), but motility in the proximal axon still appeared normal (Fig. S5, N-O). HAP1’s dynein- and dynactin-binding regions are therefore necessary for autophagosomal motility but only in the mid-axon. These findings confirm that HAP1 functions mechanistically in autophagosomal transport by activating the dynein-dynactin complex.

### JIP3 regulates autophagosomal motility primarily in the proximal axon

Since JIP1 and HAP1 are responsible for autophagosomal transport in the distal and mid-axon respectively, we next asked whether JIP3 is required for autophagosomal motility in axons, and if so, where in the axon it is functioning. We used siRNA to deplete endogenous JIP3 in neurons transfected with mScarlet-LC3; ∼70% depletion of JIP3 was achieved in PC12 cells using this approach (Fig. 7 A). In the distal and mid-axon, JIP3 depletion had at most a minor effect (Fig. 7, B-C). However, in the proximal axon JIP3 depletion induced a robust reduction in retrograde autophagosomal motility with a concomitant increase in the bidirectional/stationary fraction (Fig. 7, D-E). The loss of motility in the proximal axon could be rescued by expression of siRNA-resistant Halo-JIP3, confirming the specificity of KD (Fig. 7, D-E). However the mild effect in the mid-axon, wherein only the stationary fraction was affected, was only partially rescued by expression of siRNA-resistant JIP3 (Fig. 7 C). Thus although JIP3 is present on autophagosomes throughout the axon (Fig. 1), it primarily regulates the activity of dynein on autophagosomes in the proximal axon.

**Figure 7.**
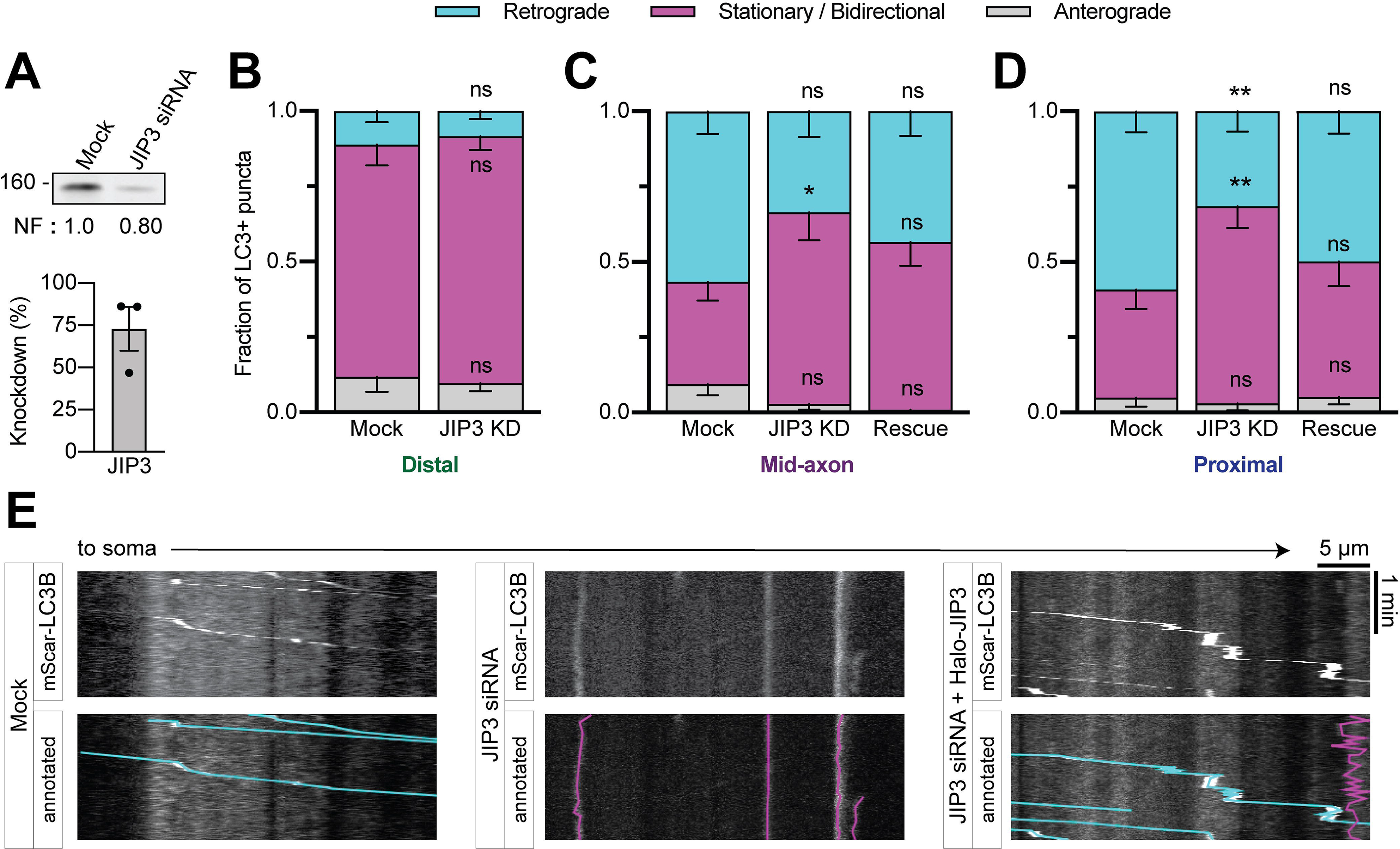
JIP3 regulates autophagosomal motility primarily in the proximal axon. **(A)** Immunoblotting and quantification of PC12 cell lysates demonstrate KD efficiency of JIP3 siRNA. *n* = 3 repeats. Normalization factor (NF) determined using Revert^TM^ Total Protein Stain. (B-D) Quantification of LC3+ puncta motile behavior in the distal **(B)**, mid-**(C)**, and proximal **(D)** axon. *n* = 12-16 neurons. Two-way ANOVA for each with Bonferroni’s multiple comparisons test (Mid-axon Retrograde Mock v. JIP3 KD, *p* = 0.1182; Mid-axon Stationary/Bidirectional Mock v. JIP3 KD, *p* = 0.0141; Mid-axon Stationary/Bidirectional Mock v. Rescue, *p* = 0.2508; Mid-axon Stationary/Bidirectional JIP3 KD v. Rescue, *p* > 0.9999; Proximal Retrograde Mock v. JIP3 KD, *p* = 0.0046; Proximal Stationary/Bidirectional Mock v. JIP3 KD, *p* = 0.0022). **(E)** Representative kymographs from the proximal axon of a mock-transfected neuron, a neuron with JIP3 siRNA, and a neuron transfected with both JIP3 siRNA and siRNA-resistant Halo-JIP3. Bars throughout figure show mean ± SEM. Symbols (ns, not significant; *, *p* < 0.05; **, *p* < 0.01) indicate Bonferroni’s test comparing to Mock.

### RILP is not a major regulator of autophagosomal motility in hippocampal axons

Our motor effector handoff mechanism implicates three main effector proteins, but it is possible that other adaptors may also contribute. In both the HAP1 (Fig. 2) and JIP3 (Fig. 7) KD experiments, a subpopulation (∼25%) of autophagosomes continued moving in the retrograde direction. This sustained motility could be due to incomplete HAP1 or JIP3 KD or may be driven by an additional dynein effector. One candidate, Rab-interacting lysosomal protein (RILP), was recently shown to be involved in autophagosomal motility in primary rat cortical neurons (Khobrekar et al., 2020). We observed RILP comigrating with autophagosomes throughout the axon (Fig. S4, A-B), and found it enriched on the autophagosomal membrane (Fig. S4, C-D) and within 40 nm of both dynein and LC3 in the axon via PLA (Fig. S4, E-H). However, when we used siRNA to deplete RILP in our primary rat hippocampal neurons (∼80% efficiency in PC12 cells; Fig. S4 I), we saw no robust KD phenotype in the distal, mid, or proximal region of the axon (Fig. S4, J-K). We did note a minor increase in the stationary fraction in the proximal axon, but this effect was not rescued by expression of siRNA-resistant Halo-RILP (Fig. S4, J-K). We therefore conclude RILP is not essential for autophagosomal motility under the conditions investigated here.

### Dynein effectors exhibit preference for autophagosomal maturation state

JIP3, HAP1, and JIP1 are active on autophagosomes in different axonal subregions. To determine what regulates the activity of these effectors, we considered how autophagosomes differ across the length of the axon. Autophagosomes mature via fusion with endolysosomal vesicles en route to the soma (Maday et al., 2012), so we asked whether dynein effectors preferentially associate with autophagosomes based on maturation state. First, we measured the maturity of autophagosomes along the axons of hippocampal neurons. We found that the majority of LC3+ puncta across the entire axon comigrated with Rab7 and LAMP1, suggesting autophagosomes fuse with late endosomes prior to exit from the distal tip (Fig. S5, A-D), consistent with previous observations made in primary mouse DRG axons (Maday et al., 2012). We next used two different markers to assess luminal acidification. Neurons were stained with LysoTracker DeepRed, a fluorescent dye that preferentially labels acidic organelles, or transfected with the dual-color fluorescent LC3 reporter, mCh-EGFP-LC3. The EGFP moiety quenches in acidic environments, resulting in a shift from dually-labeled green and red puncta to red only (Pankiv et al., 2007). Despite the early acquisition of endolysosomal membrane markers, we find autophagosomes in hippocampal axons mature somewhat slower than those in DRG axons. About 80% of autophagosomes in the mid-axon of hippocampal neurons were positive for LysoTracker (Fig. S5, E-F), compared with nearly 100% in the same region of DRG neurons (Maday et al., 2012). Further, about 60% of autophagosomes in the proximal region of hippocampal axons retained EGFP signal from their mCh-EGFP-LC3 marker (Fig. 8, A-B), compared with only about 30% of autophagosomes in proximal DRG axons (Maday et al., 2012).

**Figure 8.**
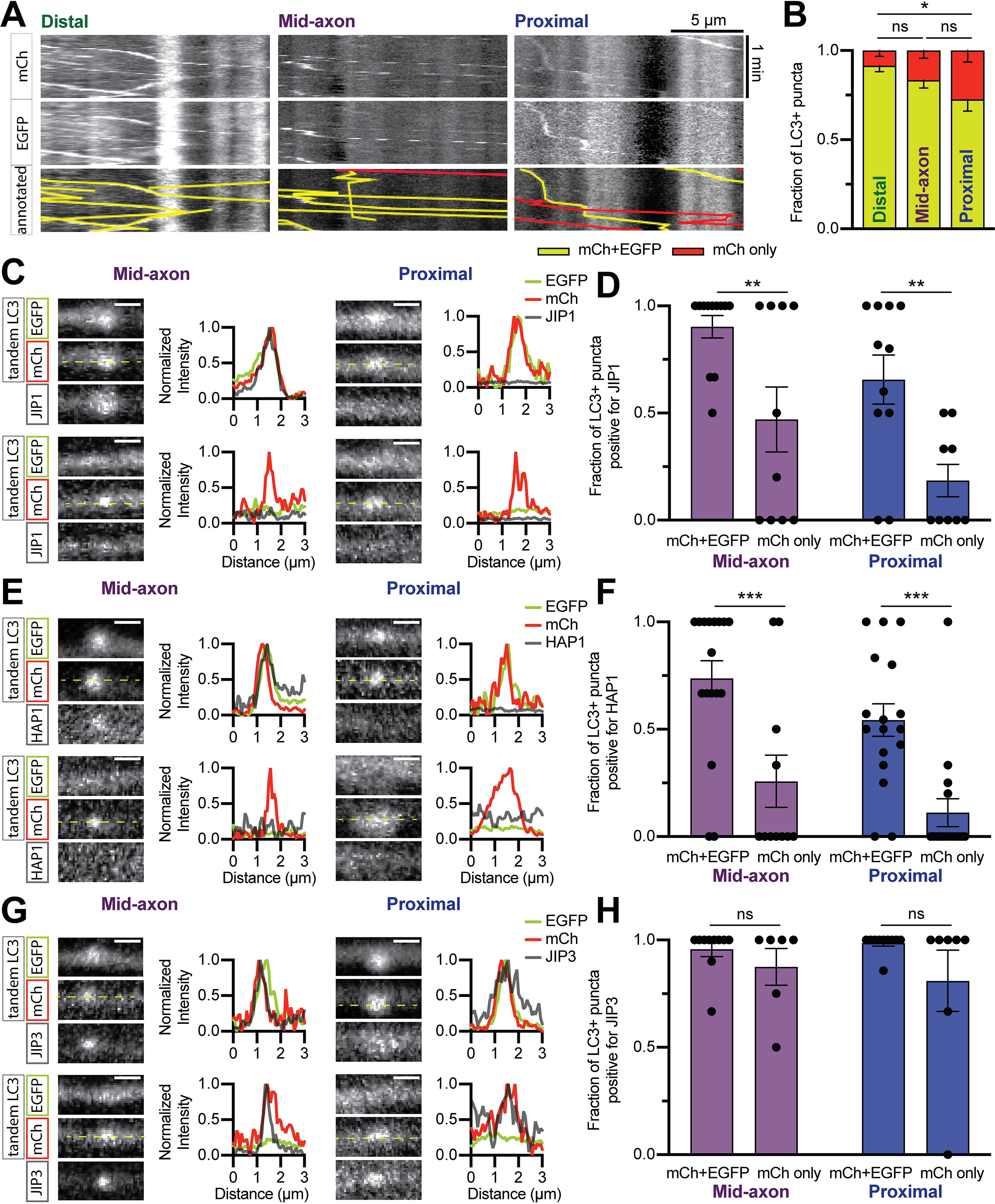
Dynein effectors exhibit preference for autophagosomal maturation state. (A-B) Example kymographs **(A)** and quantification **(B)** of the tandem mCh-EGFP-LC3 marker in different subaxonal regions. Because GFP quenches in acidic environments, autophagosomes positive for both mCh and EGFP are nonacidified and those positive only for mCh are acidified and therefore considered mature. *n* = 11-12 neurons. Two-way ANOVA with Tukey’s multiple comparisons test (Distal v. Mid-axon, *p* = 0.4452; Distal v. Proximal, *p* = 0.0195; Mid-axon v. Proximal, *p* = 0.2608). (C-H) Example micrographs, line scans, and quantifications demonstrating dynein effector association with autophagosomes of differing maturity. JIP1 **(C-D)** and HAP1 **(E-F)** significantly associate with more immature autophagosomes. JIP3 **(G-H)** associates equally with mature and immature autophagosomes. *n* = 10-18 neurons; some videos had exclusively mCh+EGFP or mCh only so the number of points on each graph may vary. Two-way ANOVA with Sidak’s multiple comparisons test (JIP1: Mid-axon mCh+EGFP v. mCh only, *p* = 0.0069; Proximal mCh+EGFP v. mCh only, *p* = 0.0092. HAP1: Mid-axon mCh+EGFP v. mCh only, *p* = 0.0007; Proximal mCh+EGFP v. mCh only, *p* = 0.0007. JIP3: Mid-axon mCh+EGFP v. mCh only, *p* = 0.6893; Proximal mCh+EGFP v. mCh only, *p* = 0.1700.) Dashed yellow lines indicate location of line scan. Scale bar, 1 µm. ns, not significant; *, *p* < 0.05; **, *p* < 0.01; ***, *p* < 0.001. Bars throughout figure show mean ± SEM.

We then tested whether dynein effectors associate preferentially with autophagosomes based on maturation state. We co-expressed mCh-EGFP-LC3 and Halo-tagged JIP1, HAP1, JIP3, or RILP then assayed comigration in the mid- and proximal axon; the distal axon was not tested because there are minimal mCh-only autolysosomes in that region (Fig. 8 B). JIP1 (Fig. 8, C-D), HAP1 (Fig. 8, E-F), and RILP (Fig. S5, G-H) were each found to associate preferentially with immature autophagosomes. In contrast, JIP3 displayed no maturation-based preference (Fig. 8, G-H), consistent with its lack of location-based preference (Fig. 1). Hence some, but not all, dynein effectors associate with autophagosomes based on their maturation state.

### Maturation state regulates the association of dynein effectors with axonal autophagosomes

We next investigated whether autophagosomal dynein scaffolding was regulated by the autophagosomal maturation process. The drug Bafilomycin A1 (BafA1) blocks autophagosomal acidification by interrupting autophagosome-lysosome fusion and blocking the vacuolar ATPase (Mauvezin et al., 2015). We treated neurons co-expressing mCh-EGFP-LC3 and Halo-HAP1 with 100 nM BafA1 or an equal volume of vehicle (DMSO) for 2 h and then imaged autophagosomes in the proximal axon. BafA1 treatment significantly increased the fraction of autophagosomes positive for EGFP, indicating an increase in unacidified autophagosomes (Fig. 9A). Accordingly, we observed an increase in the number of HAP1+ autophagosomes in the proximal axon (Fig. 9, B-C).

**Figure 9.**
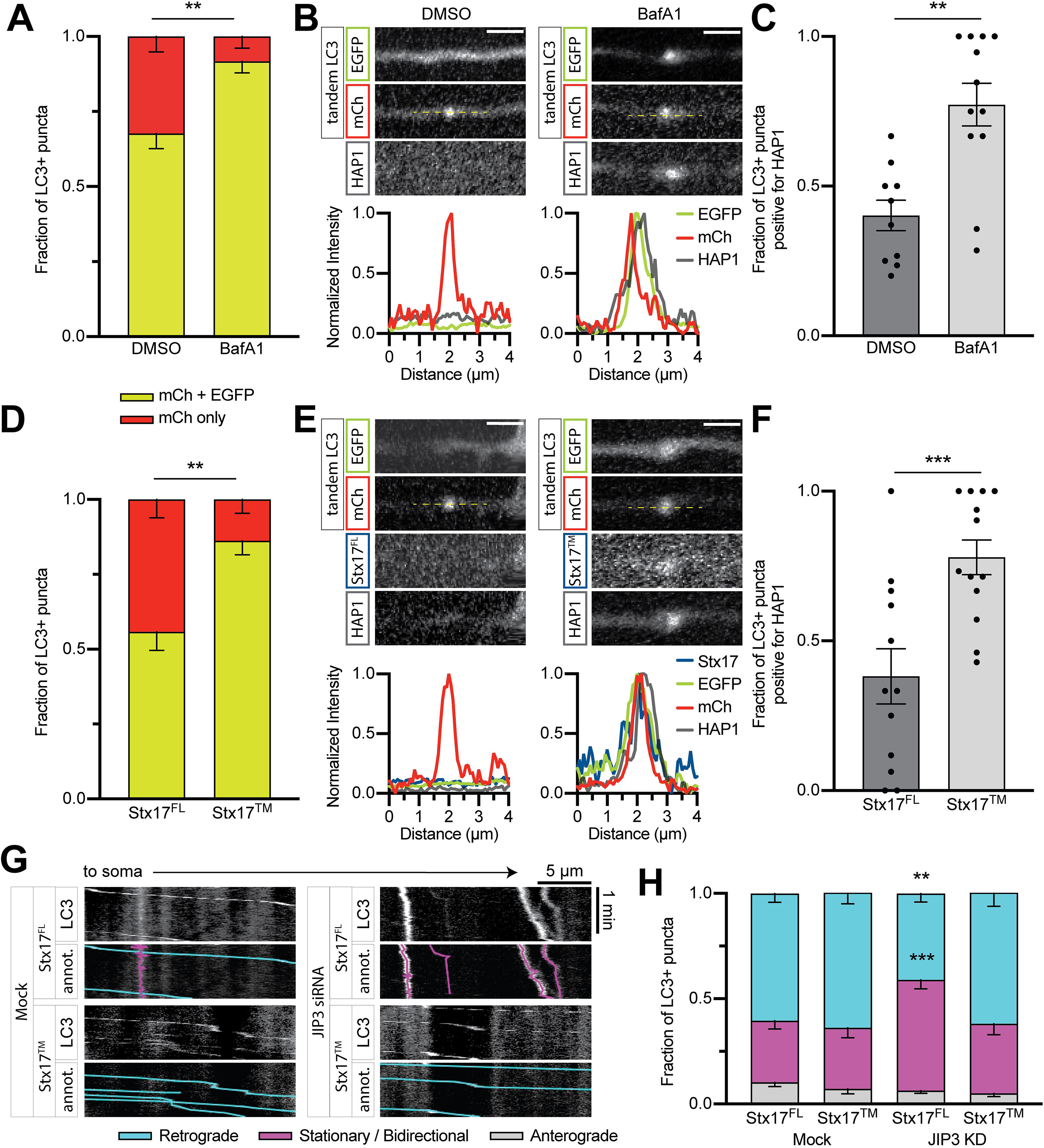
Maturation state regulates the association of dynein effectors with axonal autophagosomes. **(A)** Quantification of autophagosomal maturity using tandem mCh-GFP-LC3. BafilomycinA1 (BafA1) significantly increases the fraction of LC3+ puncta co-labeled for both mCh and EGFP above vehicle only (DMSO). *n* = 11-14 neurons. Mann-Whitney test of fraction with EGFP (*p* = 0.0050). (B-C) Example micrographs with line scans **(B)** and quantification **(C)** demonstrating HAP1 association with autophagosomes in the proximal axon is significantly higher with BafA1 treatment than with DMSO alone. *n* = 10-12 neurons. Mann-Whitney test (*p* = 0.0007). Scale bar, 2 µm. **(D)** Quantification of autophagosomal maturity using tandem mCh-GFP-LC3. Expression of the SNAP-tagged syntaxin-17 dominant negative mutant (Stx17^TM^) significantly increases the fraction of LC3+ puncta co-labeled for both mCh and EGFP above expression of SNAP-tagged full-length syntaxin-17 (Stx17^FL^). *n* = 12-13 neurons. Mann-Whitney test of fraction with EGFP (*p* = 0.0013). (E-F) Example micrographs with line scans **(E)** and quantification **(F)** demonstrating HAP1 association with autophagosomes in the proximal axon is significantly higher with Stx17^TM^ than with Stx17^FL^. *n* = 12-13 neurons. Mann-Whitney test (*p* = 0.0020). Scale bar, 2 µm. (G-H) Example kymographs **(G)** and quantification **(H)** of autophagosomal motility in the proximal axon following JIP3 siRNA transfection and/or Stx17 expression. Stx17^TM^ expression did not affect motility in the Mock condition, but Stx17^TM^ expression rescued the motility phenotype seen in JIP3 KD axons. *n* = 19-20 neurons. Two-way ANOVA with Tukey’s multiple comparisons test (Retrograde: Mock Stx17^FL^ v. Mock Stx17^TM^, *p* = 0.9372; Mock Stx17^FL^ v. KD Stx17^FL^, *p* = 0.0037; Mock Stx17^FL^ v. KD Stx17^TM^, *p* = 0.9954; KD Stx17^FL^ v. KD Stx17^TM^, *p* = 0.0019. Stationary/Bidirectional: Mock Stx17^FL^ v. Mock Stx17^TM^, *p* > 0.9999; Mock Stx17^FL^ v. KD Stx17^FL^, *p* = 0.0003; Mock Stx17^FL^ v. KD Stx17^TM^, *p* = 0.9018; KD Stx17^FL^ v. KD Stx17^TM^, *p* = 0.0044.) Symbols indicate Tukey’s test comparing to Mock. Bars throughout figure show mean ± SEM. ns, not significant; *, *p* < 0.05; **, *p* < 0.01; ***, *p* < 0.001.

To follow up on this finding, we genetically blocked autophagosome-lysosome fusion. The SNARE protein Syntaxin 17 (Stx17^FL^) interacts with SNAP29 and VAMP8 to induce autophagosome-lysosome fusion; expression of a transmembrane only (Stx17^TM^) version of the protein has a dominant negative effect on fusion (Itakura et al., 2012). We co-expressed mCh-EGFP-LC3, Halo-HAP1, and either SNAP-Stx17^FL^ or SNAP-Stx17^TM^ then imaged in the proximal axon. Expression of Stx17^TM^ increased the fraction of EGFP+ autophagosomes, indicating a shifting in the population toward a more immature state (Fig. 9 D). As a consequence, we observed more HAP1+ autophagosomes in Stx17^TM^-expressing neurons (Fig. 9, E-F).

We then tested whether the function of JIP3 on autophagosomes in the proximal axon was maturation-dependent. We co-transfected neurons with JIP3 siRNA, mCh-EGFP-LC3, and either SNAP-Stx17^FL^ or SNAP-Stx17^TM^ then measured autophagosomal transport in the proximal axon. Autophagosomes without JIP3 siRNA (Mock) displayed strong retrograde motility regardless of Stx17 construct (Fig. 9, G-H). Autophagosomes in cells co-transfected with JIP3 siRNA and Stx17^FL^ displayed a similar loss of motility (Fig. 9, G-H) as seen previously in JIP3 KD conditions (Fig. 7). However expression of Stx17^TM^ in cells transfected with JIP3 siRNA rescued normal autophagosomal motility as seen in mock conditions (Fig. 9, G-H). JIP3 therefore is primarily involved in the transport of mature autolysosomes, despite its comigration with both mature and immature autophagosomes. Hence autophagosomal maturation, and more specifically autophagosome-lysosome fusion, affects motor scaffolding on autophagosomes. Disruptions in autophagosomal transport likewise affect maturation and cargo degradation (Wong and Holzbaur, 2014), so autophagosome motility and maturity in axons are highly interdependent.

## Discussion

Here, we show that the motor effector proteins JIP1, HAP1, and JIP3 associate with autophagosomes in axons and regulate their dynein-driven motility (Fig. 10). Motile organelles, including axonal autophagosomes, associate simultaneously with both plus-end-directed kinesin motors and the minus-end-directed motor dynein (Maday et al., 2012; Hendricks et al., 2010). These opposing motors compete and/or coordinate to direct organelle motility (Kural et al., 2005; Müller et al., 2008; Hendricks et al., 2010; Kunwar et al., 2011; Hancock, 2014; Fu and Holzbaur, 2014). Regulatory proteins that modulate motor function are therefore essential to determine the net directionality of motility (Hancock, 2014; Elshenawy et al., 2019, 2020; Feng et al., 2020). For axonal autophagosomes, which demonstrate highly processive minus-end-directed motility that is essential for their function (Maday et al., 2012; Wong and Holzbaur, 2014; Fu et al., 2014), this coordinated regulation is of the utmost importance.

**Figure 10.**
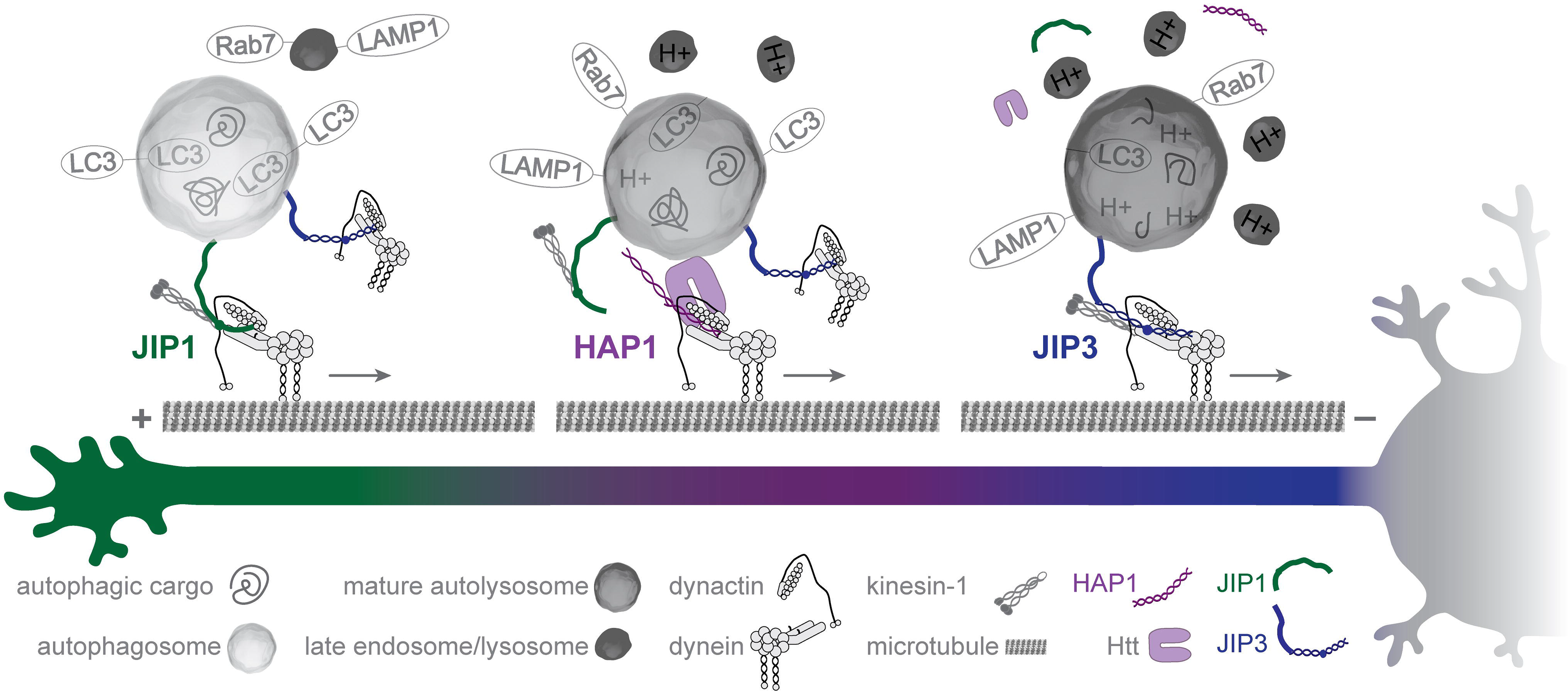
Dynein scaffolding and activation on the autophagosomal membrane is regulated by maturation along the axon. Graphic summary showing autophagosomal transport through the axon. Autophagosomes form in the distal axon (left) where they initially fuse with degradatively inactive late endosomes/lysosomes and bind JIP1, which promotes the initiation of retrograde motility by inactivating kinesin. In the mid-axon (middle), HAP1 and its interacting partner Htt are responsible for scaffolding the dynein-autophagosome relationship, with HAP1 directly activating the dynein complex via canonical and novel interaction regions for dynein-dynactin. Finally, as the autophagosome continues to mature via fusion with more late endosomes/lysosomes, HAP1 and JIP1 disassociate and JIP3 promotes the continued retrograde motility of the autolysosome in the proximal axon (right). The autophagosome then arrives at the soma, where competent lysosomes are enriched and degraded materials can be recycled.

Previous work has implicated motor-binding proteins JIP1, HAP1, and JIP3 in autophagosomal transport (Fu et al., 2014; Wong and Holzbaur, 2014; Hill et al., 2019), but the interplay between them was not understood. Our work demonstrates these proteins drive autophagosomal transport in discrete subaxonal regions. JIP1 specifically interacts with immature autophagosomes and dynein distant from the soma. HAP1 regulates the motility of immature autophagosomes in the mid-axon and dissociates from mature autolysosomes in the proximal axon. JIP3 binds autophagosomes indiscriminately, but primarily drives the motility of autolysosomes in the proximal axon. This handoff between dynein regulators on a single cargo has not been observed before but may represent a more general paradigm that is relevant in the transport of other organelles, especially those that mature (e.g. phagosomes) and/or traverse long distances (e.g. signaling endosomes). For example, previous work has shown that dynein effector Hook1 drives the retrograde motility of BDNF-containing signaling endosomes, but only in the distal axon (Olenick et al., 2019). The work described here suggests a different dynein effector may take over in the mid- and proximal axon.

We find JIP1 localizes specifically to autophagosomes in the distal and mid-axon (Fig. 1). This is consistent with previous work showing JIP1 is required for efficient exit from the distal end of the axon and continued inhibition of kinesin in the mid-axon (Fu et al., 2014). JIP1 interacts with both kinesin and dynein, and phosphorylation of residue S421 acts as a switch between anterograde and retrograde transport (Fu and Holzbaur, 2013). This molecular switch is required for the initiation of autophagosomal transport in the distal axon (Fu et al., 2014). Further, JIP1 binds LC3 directly and LC3-binding blocks KHC activation in single molecule motility assays (Fu et al., 2014). It is not yet known whether JIP1 directly activates dynein in addition to inactivating kinesin.

Our work is consistent with the previous finding that HAP1 and Htt are required for axonal autophagosome transport in DRG axons (Wong and Holzbaur, 2014), and extends this observation by showing that HAP1 and Htt are specifically required in the mid-axon (Fig. 2). We identify conserved dynein- and dynactin-binding sites within HAP1 (Fig. 3-4) that are necessary for autophagosomal motility (Fig. 6). These interactions, together with HAP1’s ability to induce dynein activity in single molecule motility assays (Fig. 5), introduce HAP1 to the growing family of dynein activating adaptors (Reck-Peterson et al., 2018; Olenick and Holzbaur, 2019).

The CC1 box motif forms a hydrophobic pocket for the binding of dynein subunit LIC1 (Lee et al., 2020). This same hydrophobic pocket is formed in multiple unrelated dynein effector families: the CC1-box-containing family, the Hook-containing family, and the EF-hand-containing family (Lee et al., 2018; Celestino et al., 2019; Gama et al., 2017; Lee et al., 2020). Our work shows HAP1’s CC1 box is required for dynein activation on autophagosomes in neurons (Fig. 6). The CC1 box is likewise necessary for dynein activator function in vivo: a de novo mutation in the CC1 box of Drosophila BICD causes a hypomorphic loss-of-function phenotype (Oh 2000). The Spindly motif, a short sequence conserved amongst many dynein complex activators, binds the pointed end complex of dynactin’s Arp1 filament (Gama et al., 2017). This motif is required for robust Hook3-dynein-dynactin motility (Schroeder and Vale, 2016) and for the binding of Spindly protein to dynactin (Gama et al., 2017). A popular hypothesis suggests the pointed-end complex protein p25 contains a mutually exclusive binding site that either binds an activator’s Spindly motif or the tail of auto-inhibited p150^Glued^; therefore activator binding to p25 releases p150^Glued^ from autoinhibition (Cianfrocco et al., 2015; Schroeder and Vale, 2016; Qiu et al., 2018; Lau et al., 2020). Our co-immunoprecipitation experiments (Fig. 3) and axonal transport studies (Fig. 6) confirm that HAP1 contains a functional Spindly motif and therefore contributes to dynactin disinhibition.

We also identify a third conserved binding site through which HAP1 binds dynactin, independent of the Spindly motif. This p150^Glued^ -binding site is conserved among the HAP/TRAK and HOOK families of dynein effectors but is not found in other dynein effectors such as BICD2 (Fig. S3). Yeast two-hybrid assays showed this region binds the second coiled coil region of p150^Glued^ (Engelender et al., 1997), which has been suggested to regulate membrane binding by dynactin (Kumar et al., 2001). We show here that a motif comprising HAP1 residues 317-334 is required for p150^Glued^ binding and point mutations in this motif can disrupt axonal transport of both HAP1 itself (Fig. 4) and autophagosomes (Fig. 6). The function of this *Glued* binding motif remains to be tested in other proteins, and structural studies will be required to better understand the interaction and its implications for the dynein-dynactin complex.

We find that Htt is required in complex with HAP1 to direct mid-axon autophagosomal motility (Fig. S2). We hypothesize the primary role of Htt in autophagosomal transport is as a scaffold to bring the dynein-dynactin-HAP1 complex into contact with the autophagosome. Htt interacts with HAP1 (Li et al., 1996), dynein (Caviston et al., 2007), and dynactin (Engelender et al., 1997; Li et al., 1998), and both the dynein-binding and HAP1-binding regions of Htt are required for autophagosomal transport (Wong and Holzbaur, 2014). Further, Htt interacts with multiple autophagosomal membrane proteins including LC3B and GABARAPL1 (Ochaba et al., 2014), making it a perfect scaffold for autophagosomal motility (Saudou and Humbert, 2016).

We find that JIP3 is present on autophagosomes throughout the axon, but it primarily drives the motility of mature autolysosomes in the proximal axon (Fig. 7). JIP3 interacts with both kinesin and dynein-dynactin (Cockburn et al., 2018; Cavalli et al., 2005; Vilela et al., 2019) and activates kinesin-dependent motility in vitro (Sun et al., 2011). JIP3 regulates axonal autophagosome transport in *C. elegans* (Hill et al., 2019), and our work demonstrates a similar role in vertebrates. Previous studies have implicated JIP3 in the motility of endolysosomes (Drerup and Nechiporuk, 2013; Gowrishankar et al., 2017; Brown et al., 2009); notably mature autolysosomes, on which JIP3 is most active, share many of their membrane proteins with endolysosomes. While there is a clear genetic interaction between autophagosomal maturation and JIP3-dependent transport (Hill et al., 2019), more work is necessary to understand how JIP3 is regulated on autophagosomes.

In both the HAP1 (Fig. 2) and JIP3 (Fig. 7) knockdown experiments, we observed low levels of persistent retrograde transport due either to incomplete knockdown or other motor effector proteins. RILP was recently shown to mediate autophagosomal transport in rat cortical neurons (Khobrekar et al., 2020), so we tested RILP in our system. While we saw comigration of RILP with LC3+ puncta, RILP KD did not induce a robust effect on autophagosomal transport in rat hippocampal neurons. One possible explanation for the discrepancy may be that Khobrekar et al. were studying neurons at an earlier developmental timepoint. Other motor regulators (Cai et al., 2010; Cheng et al., 2015) may also be involved in autophagosomal transport in axons.

We find here that maturation regulates the association and function of dynein effectors on the autophagosome. HAP1 is abberantly retained on proximal autophagosomes following pharmacological or genetic inhibition of autophagosome maturation, indicating that maturation induces dissociation of some dynein effectors (Fig. 9). Additionally, JIP3 interacts indiscriminately with autophagosomes but primarily regulates the transport of mature autolysosomes (Fig. 7 and 9). Maturation could affect motor scaffolding through a number of mechanisms. The dissociation of effectors associated with immature autophagosomes (HAP1, JIP1) may be caused by loss of binding partners (e.g. LC3) on the fused membrane. Or, a lysosomal membrane protein acquired during autophagosome-lysosome fusion could promote dissociation of immature dynein scaffolds and/or activate mature dynein scaffolds (JIP3). Alternatively, acidification of the mature autolysosome could trigger an inside-out signaling cascade causing dissociation and/or activation of scaffolding proteins. Further research is necessary to tease out these possibilities.

Since both axonal transport and autophagy are strongly associated with neuronal health and neurodegeneration, future studies on the regulation of axonal autophagosome transport are likely to provide valuable insights into human disease (De Pace et al., 2018). Htt, HAP1, JIP1, and JIP3 have all been implicated in neurodegenerative disease (Waeber et al., 2000; Helbecque et al., 2003; Gunawardena et al., 2003; Morfini et al., 2009; Beeler et al., 2009; Fu and Holzbaur, 2013; White et al., 2015; Weiss and Littleton, 2016; Choudhary et al., 2017; Gowrishankar et al., 2017). Further research into axonal transport and autophagy may guide the development of effective therapies for neurodegeneration.

## Materials and Methods

### Plasmids and reagents

Constructs, all of which were verified by DNA sequencing, include the following:

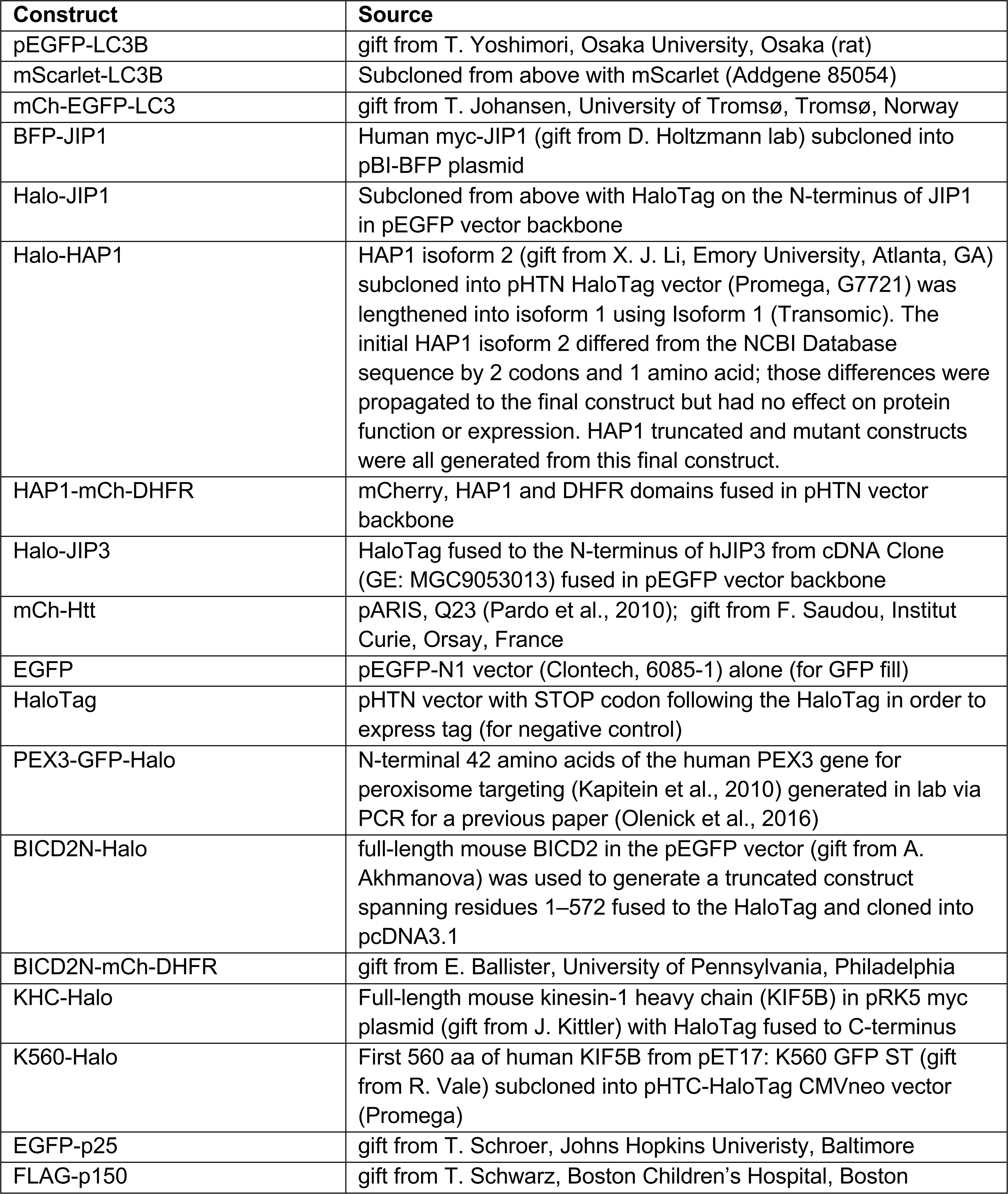
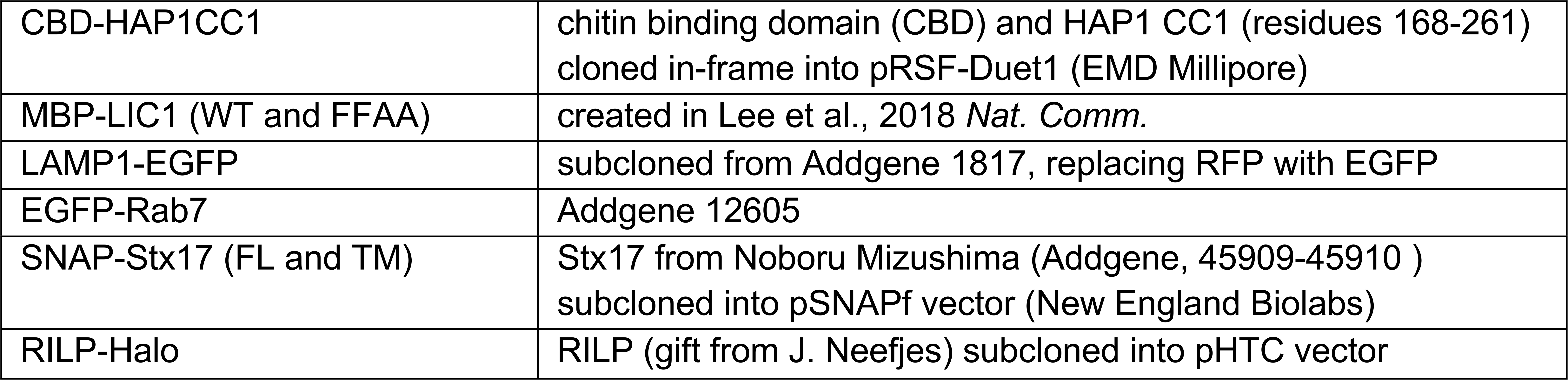

As an alternative to transfecting an LC3 plasmid, some experiments used the Premo^TM^ Autophagy Sensor BacMaM 2.0 LC3B-GFP (ThermoFischer, P36235) which introduces DNA via insect Baculovirus with a Mammalian promoter. Individual ON-TARGETplus siRNA to HAP1 (5’-GAAGUAUGUCCUCCAGCAAUU-3’), Htt (5’-GCAGCUUGUCCAGGUUUAUUU), JIP3 (5’-CAGCUGGCUUUAGCCAGCGUCGCAAUU), and RILP (5’-CGGUGAACAUCUUGGUCUG) were obtained from Dharmacon (Horizon Discovery). Chemical compounds used include LysoTracker Deep Red (ThermoFisher, L12492) and BafilomycinA1 (SigmaAldrich, B1793). HaloTag constructs were labeled with Janelia Fluor 646 HaloTag (Promega, GA1120), Janelia Fluor 549 (Promega, GA1110), or TMR (Promega, G8251). SNAP-Tag constructs were labeled with SNAP-Cell® 430 (New England BioLabs, S9109S) or SNAP-Cell® 647-SiR (New England BioLabs, S9102S).

Antibodies include the following:

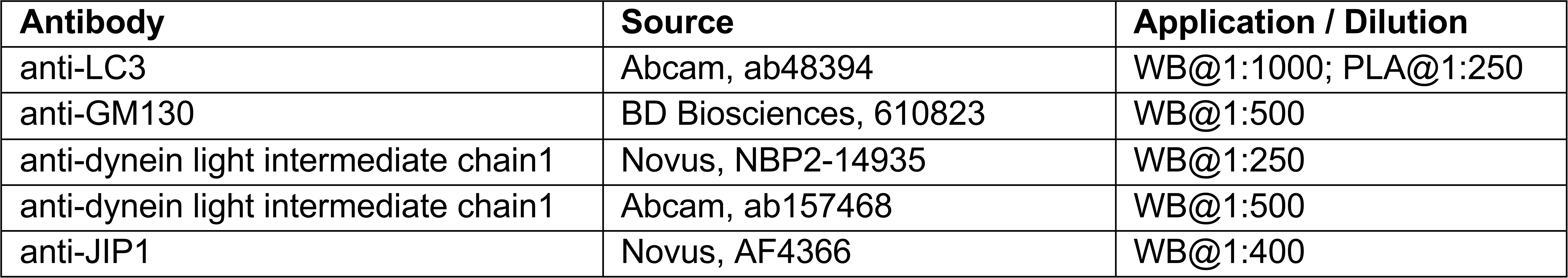
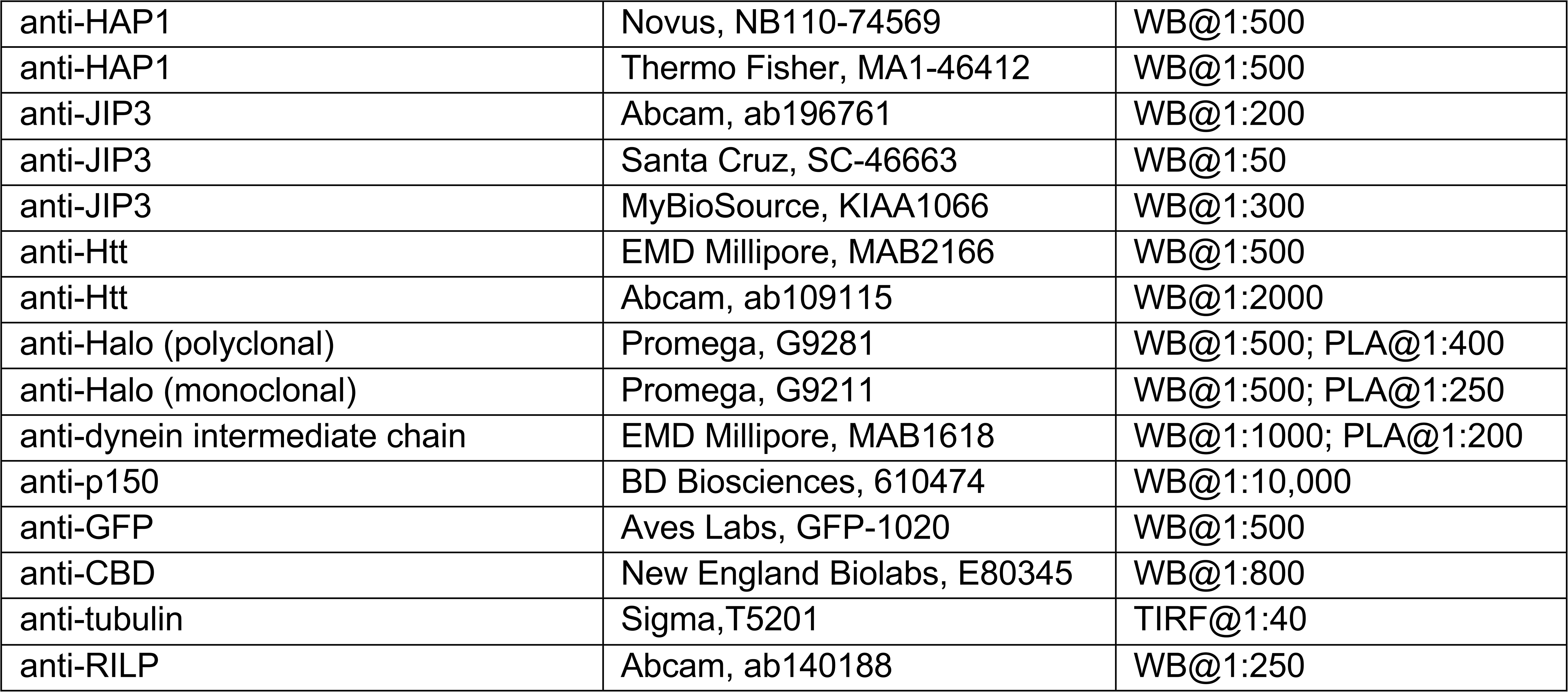

### Primary hippocampal culture

Sprague Dawley rat hippocampal neurons at embryonic day 18 were obtained from the Neurons R Us Culture Service Center at the University of Pennsylvania. Cells (proximity ligation assay, 40,000 cells on 7mm glass; live imaging, 200,000 cells on 20 mm glass) were plated in glass-bottom 35 mm dishes (MatTek) that were precoated with 0.5 mg/ml poly-L-lysine (Sigma Aldrich). Cells were initially plated in Attachment Media (MEM supplemented with 10% horse serum, 33 mM D-glucose, and 1 mM sodium pyruvate) which was replaced with Maintenance Media (Neurobasal [Gibco] supplemented with 33 mM D-glucose, 2 mM GlutaMAX (Invitrogen), 100 units/ml penicillin, 100 mg/ml streptomycin, and 2% B-27 [ThermoFisher]) after 5-20 h. Neurons were maintained at 37 C in a 5% CO2 incubator; AraC (final conc. 1 µM) was added the day after plating to prevent glia cell proliferation. For DNA only or DNA + siRNA transfections, neurons (4-6 DIV) were transfected with 0.35–1.5 µg of total plasmid DNA and optional 45 pmol siRNA using Lipofectamine 2000 Transfection Reagent (ThermoFisher, 11668030) and incubated for 18-48 h. For siRNA only transfections (Fig. 2 & S2), neurons were transfected with 45 pmol siRNA using Lipofectamine RNAiMAX (ThermoFisher, 13778030) and incubated for 36–48 h; to label LC3 in those experiments, ∼ 20 MOI BacMAM LC3 (40 uL) was added immediately following transfection.

### Live-cell neuron imaging and analysis

One hour prior to imaging, HaloTag® ligands (TMR, Promega, G8251; Janelia Fluor® 549, Promega GA1110; Janelia Fluor® 646, Promega, GA1120) and/or SNAP-tag® ligands (Janelia Fluor® 646, provided by Luke Lavis, Janelia; SNAP-Cell® 647-SiR, New England Biolabs, S9102S) were applied for 15 min at a final concentration of 100 nM, followed by a 30-45 min washout; SNAP-tag® ligand Blue 430 (New England Biolabs, S9109S) was applied for 30 min at a final concentration of 2 µM, followed by a 30 min washout. In applicable experiments, neurons were incubated with Lysotracker (25 nM) for 15-30 min, which was then removed for imaging. In applicable experiments, BafilomycinA1 (100 nM) or DMSO was added 2h prior to imaging, and then neurons were imaged in the third hour of continued treatment. Neurons were imaged in Imaging Media (HibernateE [Brain Bits] supplemented with 2% B27 and 33 mM D-glucose). Autophagosome behavior was monitored in the proximal (<100 µm from the soma), distal (<100 µm from the distal tip), or mid-axon of 6–8 DIV neurons imaged at a rate of 1 timepoints/sec for 2-3 min. Neurons were imaged in an environmental chamber at 37°C on a Perkin Elmer UltraView Vox spinning disk confocal on a Nikon Eclipse Ti Microscope with an Apochromat 100 x 1.49 numerical aperture (NA) oil-immersion objective and a Hamamatsu EMCCD C9100-50 camera driven by Volocity (PerkinElmer).

Kymographs were generated in ImageJ (https://imagej.net/ImageJ2) using the MultiKymograph plugin (line width, 5) and analyzed either in ImageJ or using the MatLab program KymoSuite (J. Nirschl, University of Pennsylvania). Puncta were classified as either anterograde (moving ≥10µm towards the axon tip), retrograde (moving ≥10µm towards the soma), or Stationary/bidirectional (net movement <10µm during the video). Because fluorescent LC3 is cytosolic (as well as punctate) and neurites occasionally crossed in culture, raw videos were referenced throughout kymograph analysis to ensure only real puncta (≥ 1.5 SD from the axon mean) were included in analyses. All comigration analyses were performed using kymographs. Line scans were generated for presentation purposes from raw video stills and normalized either within that line (for positive channels) or to the local region (for negative channels; surrounding ∼10µm area).

### Autophagosome fractionation

Enriched autophagosome fractions were isolated by modifying a published protocol (Strømhaug et al., 1998) and detailed protocols and validations can be found in Goldsmith et al. (Manuscript in preparation). Briefly, brains were collected from wildtype or GFP-LC3B transgenic mice on the C57BL/6J background (Ref 14699058) and homogenized in a tissue grinder in an ice cold buffered 10mM Hepes, 1mM EDTA, 250 mM sucrose solution, then subjected to three differential centrifugations through Nycodenz and Percoll discontinuous gradients to isolate vesicles of the appropriate size and density. The autophagosome enriched fraction was then divided and either immediately lysed for the identification of all internal and externally-associated proteins on autophagosomes (A fraction), treated with 10 μg proteinase-K for 45min at 37°C to degrade externally associated proteins and enrich for membrane-protected autophagosome cargo (P fraction), or membrane permeabilized by the addition of 0.2% triton x-100 prior to proteinase K treatment to confirm proteinase K efficacy (T fraction). The lysis buffer used contained a final concentration of 0.5% NP-40 with 1x protease and phosphatase inhibitors, PMSF and Pepstatin A. Protein concentration was measured by Bradford assay and equal amounts of protein in denaturing buffer were run on SDS-PAGE gels.

### Immunoblotting

For fluorescence Western blotting, samples were analyzed by SDS-PAGE and transferred onto PDVF Immobilon FL (Millipore). Membranes were dried for at least 1 h, rehydrated in methanol, and stained for total protein (LI-COR REVERT Total Protein Stain). Following imaging of the total protein, membranes were destained, blocked for 1 h in TrueBlack® Blocking Buffer (Biotium, 23013), and incubated overnight at 4°C primary with antibodies diluted in TrueBlack® Antibody Dilutent (Biotium, 23013) with 0.2% Tween-20. Membranes were washed four times for 5 min in 1xTBS Washing Solution (50 mM Tris-HCl pH 7.4, 274 mM NaCl, 9 mM KCl, 0.1% Tween-20), incubated in secondary antibodies diluted in TrueBlack® Antibody Dilutent (Biotium, 23013) with 0.2% Tween-20 and 0.01% SDS for 1 hr, and again washed four times for 5 min in the washing solution. Membranes were immediately imaged using an Odyssey CLx Infrared Imaging System (LI-COR). In a limited number of cases, the membrane was stripped using NewBlot^TM^ IR Stripping Buffer (LI-COR, 928-40028) according to manufacturer’s instructions. Band intensity was measured in the LI-COR Image Studio application.

### Proximity ligation assay (PLA)

Neurons were transfected (Lipofectamine 2000) with 0.3 µg EGFP plasmid (for GFP fill) and 0.5 µg Halo-tagged effector following above protocol then 24 h later (DIV 7–8) fixed in PBS containing 4% paraformaldehyde and 4% sucrose for 8 min. Duolink^TM^ In Situ PLA Mouse/Rabbit kit with red detection reagents (Sigma-Aldrich, DUO92101-1KT) was used according to manufacturer’s protocol. A Halo antibody (either Mouse G9211 or Rabbit G9281) was used in every experiment along with either an LC3 antibody (Rabbit ab48394), a dynein intermediate chain antibody (Mouse MAB1618), or no second 1° antibody (negative control). Both 2° antibodies (Mouse and Rabbit) were added for all experiments (including negative control). Z-stacks (0.25 µm steps) were acquired on an inverted epifluorescence microscope (DMI6000B; Leica) with an Apochromat 63 x 1.4 NA oil-immersion objective and a charge-coupled device camera (ORCA-R2; Hamamatsu Photonics) using LAS-AF software (Leica). Puncta were counted manually using ImageJ.

### Cell line culture

COS-7 (ATCC) cells were maintained in DMEM (Corning) supplemented with 1% GlutaMAX and 10% FBS. PC12 (ATCC) cells were maintained in DMEM supplemented with 1% GlutaMAX, 5% FBS, and 5% horse serum. Cells were maintained at 37 C in a 5% CO2 incubator. For PRA experiments, COS-7 cells were plated on 35 mm glass-bottom plates; 24h prior to imaging cells were co-transfected with human PEX3^1-42^-GFP-Halo and either human HAP1-mCherry-eDHFR or mouse BICD2^1-572^-mCherry-eDHFR using FuGENE 6 (Promega; 1 µg total DNA) and 48h prior to imaging with control siRNA using Lipofectamine RNAiMAX (Thermo Fisher). For motility assays and co-immunoprecipitation experiments, COS-7 cells were plated on 10 cm plates and transfected 24h prior to lysis using FuGENE 6 (Promega; 6-12 µg total DNA). For siRNA tests, PC12 cells were plated in 6 well dishes and transfected 48h prior to lysis with 45 pmol siRNA using Lipofectamine RNAiMAX (Thermo Fisher); cells were lysed in RIPA buffer (50 mM Tris-HCl pH 7.4, 150 mM NaCl, 0.1% Triton X-100, 0.5% deoxycholate, 0.1% SDS, and protease inhibitors (1 mM PMSF, 0.01 mg/ml Nα-p-tosyl-L-arginine methyl ester [TAME], 0.01 mg/ml leupeptin, 0.001 mg/ml pepstatin A, 1 mM DTT) for 30 min at 4 C followed by centrifugation at 4 C for 10 min at 17,000 x g and a BCA assay of the supernatant to determine total protein concentration. Cells were routinely tested for mycoplasma using a MycoAlert detection kit (Lonza, LT07). COS7 and PC12 cells were authenticated by ATCC.

### Protein purification and pull-down

BICD2N-Halo and MBP-LIC1 (WT and FFAA) were purified as described previously (Lee et al., 2018 *Nat. Comm.*). CBD-HAP1CC1 was expressed in ArcticExpress(DE3) RIL cells (Agilent Technologies), grown in Terrific Broth (TB) medium for 6 hours at 37°C to a density of ∼1.5 to 2 optical density at 600 nm (OD600), followed by 24 hours at 10°C in the presence of 0.4 mM isopropyl-β-d-thiogalactoside (IPTG). Cells were harvested by centrifugation, resuspended in 20 mM Hepes (pH 7.5), 200 mM NaCl, 10 mM imidazole, and 2 mM PMSF and lysed using a microfluidizer (Microfluidics). The protein was first purified using Ni-NTA resin according to the manufacturer’s protocol and eluted using the same buffer supplemented with 300 mM imidazole. A final step of purification was performed on a gel filtration SD200HL 16/60 column (GE Healthcare) in 20 mM Hepes (pH 7.5) and 100 mM NaCl.

Pull down experiments were performed in 50 mM Hepes (pH 7.5) with 100 mM NaCl, 1 mM DTT, and 0.1% Tween20. MBP-LIC1 (50 pmol) and either BICD2N-Halo (50 pmol) or CBD-HAP1CC1 (100 pmol) was incubated with resin (Magne® HaloTag, Promega, G7281; Chitin, New England BioLabs, S6651; or Amylose, New England BioLabs, E8021) for 1 h at 4°C. Following incubation, resin was washed 3 x 500 µL in pulldown buffer, then resuspended in 100 µL denaturing buffer and boiled to release the bound proteins. Protein pulldown was analyzed by Western Blot.

### Co-immunoprecipitation

Dynabeads Protein G (Thermo Fisher 10007D) were incubated with anti-p150 antibody (BD 610474) for 15 min prior to addition of lysate. COS-7 cells were lysed 24 h after transfection in 50 mM Hepes (pH 7.4), 1 mM EDTA, 1 mM MgCl2, 25 mM NaCl, with 0.5% Triton-X and protease inhibitors (1 mM PMSF, 0.01 mg/ml TAME, 0.01 mg/ml leupeptin, 0.001 mg/ml pepstatin A, 1 mM DTT) and then clarified at 17 x g at 4 °C for 10 min. Dynabeads or Magne® HaloTag beads were then washed with lysis buffer and incubated with lysate for 15 min at 25°C. Following incubation, beads were washed 3 x 300 µL in lysis buffer, then resuspended in 60 µL denaturing buffer and boiled to release the bound proteins. Co-immunoprecipitation was analyzed by Western Blot.

### Peroxisome recruitment assay (PRA)

The uncaged dimerizer TMP-Htag (Ballister et al., 2015) was dissolved in DMSO to 10 mM concentration and stored in at -20°C. After capturing stills and a 2 min pre-dimerizer video, the dimerizer (or DMSO) was applied at a working concentration of 10 μM dropwise directly to the cells followed by a 25m video of the dimerization (0.5 frames/s) followed by post-dimerizer stills. Imaging medium was L-15 medium (Thermo Fisher) supplemented with 10% fetal bovine serum. Videos were acquired on an inverted epifluorescence microscope (DMI6000B; Leica) with an Apochromat 63 x 1.4 NA oil-immersion objective and a charge-coupled device camera (ORCA-R2; Hamamatsu Photonics) using LAS-AF software (Leica). Color-coded time projections were generated in ImageJ using a publicly available plugin (K. Miura, EMBL Heidelberg, https://imagej.net/Temporal-Color_Code).

### Motility assay

The movement of HAP1- or BICD2N-containing complexes from cell extracts was tracked using TIRF microscopy. Motility assays were performed in flow chambers constructed with a glass slide and a coverslip silanized with PlusOne Repel-Silane ES (GE Healthcare), held together with vacuum grease to form a ∼10 μl chamber. Rigor kinesin-1_E236A_ (0.5 µM) was non-specifically absorbed to the coverslip (Wagenbach et al., 2008) and the chamber was then blocked with 5% pluronic F-127 (Sigma-Aldrich). 250 nM GMPCPP microtubule seeds, labeled at a 1:40 ratio with HiLyte Fluor 647 tubulin (Cytoskeleton, Denver, CO), were flowed into the chamber and immobilized by interaction with rigor kinesin-1_E236A_. 11.25 µM free tubulin (labeled at a 1:20 ratio with HiLyte Fluor 488 tubulin) was added with the lysate to grow dynamic microtubules from the seeds. COS-7 cells grown in 10 cm plates to 70–80% confluence expressing full-length Halo-tagged HAP1, BICD2N or HaloTag alone were labeled with TMR-HaloTag ligand (Promega, Madison, WI) 18–24 h post-transfection then lysed in 100 μl lysis buffer [40 mM Hepes (pH 7.4), 120 mM NaCl, 1 mM EDTA, 1 mM ATP, 0.1% Triton X-100, 1 mM PMSF, 0.01 mg ml^-1^ TAME, 0.01 mg ml^-1^ leupeptin, and 1 μg ml^-1^ pepstatin-A]. Cell lysates were clarified by centrifugation (17,000*g*) and diluted in P12 motility buffer [12 mM Pipes (pH 6.8), 1 mM EGTA, and 2 mM MgCl_2_] supplemented with 1 mM Mg-ATP, 1 mM GTP, 0.08 mg ml^-1^ casein, 0.08 mg ml^-1^ bovine serum albumin, 2.55 mM DTT, 0.05% methylcellulose, and an oxygen scavenging system (0.5 mg ml^-1^ glucose oxidase, 470 U ml^-1^ catalase, and 3.8 mg ml^-1^ glucose). All the videos (2 min, 4 frames s-1) were acquired at 37 °C using a Nikon TIRF microscopy system (Perkin Elmer, Waltham, MA) on an inverted Ti microscope equipped with a 100× objective and an ImageEM C9100-13 camera (Hamamatsu Photonics, Hamamatsu, Japan) with a pixel size of 0.158 µm and controlled with the program Volocity (Improvision, Coventry, England). At least 5 microtubules per video were analyzed by generating kymographs using the MultiKymograph plugin of ImageJ and analyzed in Excel (Microsoft, Redmond, WA). During acquisition, the seeds (647) were imaged at a rate of 1 frame min^-1^ and the free tubulin (488) at 12 frames min^-1^ and only non-bundled microtubules with a clear plus-end (one end clearly growing faster and longer away from the seed) were analyzed. At least 15 microtubules were analyzed per replicate; 3 biological and technical replicates were performed for a final n = 57 microtubules per condition.

Processive runs were defined as events (≥1 SD above the background) ≥ 1 µm in length and lasting ≥ 1 s. Runs ending within 0.5 µm of the microtubule minus end were excluded from the run length analysis. Statistical analyses were performed in Prism (GraphPad, San Diego, CA). A two-tailed t-test was used for velocity analysis, a two-tailed Mann-Whitney test was used for run-length analysis, and a Kruskal-Wallis test followed by a Dunn’s multiple comparisons test was used for analysis of the number of events.

### Kinesin microtubule binding assay

COS-7 cells grown in 10 cm plates to 70–80% confluence expressing Halo-tagged K560 or KIF5B (KHC) were labeled with TMR-HaloTag ligand (Promega, Madison, WI) 18–24 h post-transfection then lysed in P12 buffer (as described above) supplemented with 20 µM Taxol, 1 mM Mg-ATP, 0.1% Triton, and 10 µM DTT supplemented with protease inhibitors (described above). Separate plates expressing Halo-HAP1 or Halo-HAP1 + mCh-Htt were likewise lysed but not labeled. All lysates were clarified by a low speed (17000 × *g*) and a high speed (287582 × *g*) centrifugation. Flow chambers were assembled as described above. A 1:40 dilution of monoclonal anti-tubulin antibody was flowed in and incubated 5 minutes. The chamber was then blocked with two 5 minute incubations of 5% pluronic F-127 (Sigma). Labeled (labeling ratio of 1:40, HiLyte 488, Cytoskeleton) Taxol-stabilized microtubules were then flowed into the chamber and immobilized on the antibody. Finally, diluted cell lysates were flowed in with P12 buffer containing 10 mM AMPPNP, 20 µM Taxol, 0.3 mg/ml bovine serum albumin, 0.3 mg/ml casein, 10 mM DTT, and an oxygen-scavenging system (described above). One three minute video was acquired for each chamber at 0.067 frames/s. The mean fluorescence intensity for 5 unbundled microtubules per video was collected at 30 second intervals in ImageJ.

### Microtubule pelleting assay

Unlabeled tubulin was polymerized at 5 mg/ml in BRB80 (80 mM Pipes, 1 mM EGTA, and 1 mM MgCl2 (pH 6.8)) with 1 mM GMPCPP. To generate cell lysate, COS7 cells transfected with Halo-HAP1 (for 18–20 h) were lysed in BRB80 buffer with 0.5% Triton X-100 and protease inhibitors (as described above) and clarified with two centrifugation steps (at 17,000 and 27,000 x g). 0 or 5 µM microtubules were incubated with an equal concentration of cell lysate at 37°C for 20 min, with 1 mM Mg-ATP or 150 mM KCl where appropriate. Then samples were centrifuged at 10,000 x g at 25 °C for 20 min. The supernatant and the pellet were then separated, denatured, and analyzed by SDS-PAGE. Protein in the pellet and supernatant fractions was analyzed by Western Blot.

## Acknowledgements

This research was supported by NIH grants NIH grants R35 GM126950 to E.L.F.H. and RM1 GM136511 to R.D. and E.L.F.H. S.E.C. was a recipient of an NSF Graduate Research Fellowship (DGE-1845298) and P.J.C. was supported by an NIH T32 grant (AR053461). The authors declare no competing financial interests. We thank Mariko Tokito for technical assistance; Chanat Aonbangkhen for the dimerizer reagent; and Andrea Stavoe, Chantell Evans, Alex Boecker, and Adam Fenton for insights and discussions.

## Author contributions

Sydney E. Cason, Conceptualization, Resources, Data curation, Formal analysis, Validation, Investigation, Visualization, Methodology, Project administration, Writing— original draft and review/editing; Peter Carman, Methodology, Resources, Writing— review/editing; Claire Van Duyne, Investigation, Formal analysis, Writing—review/editing; Juliet Goldsmith, Resources, Validation, Writing—review/editing; Roberto Dominguez, Conceptualization, Supervision, Writing—review/editing; Erika L.F. Holzbaur, Conceptualization, Supervision, Funding acquisition, Project administration, Writing— original draft and review/editing

## Suppemental Material

Fig. S1 shows candidate effectors closely apposed to dynein and autophagosomes in axons. Fig. S2 compares Htt KD in the different axonal subregions and illustrates the conservation of the novel *Glued* binding motif. Fig. S3 shows HAP1 cannot activate kinesin in vitro and its dynein activation is not required on autophagosomes in the proximal axon. Fig. S4 illustrates RILP associates with axonal autophagosomes but does not play a role in their transport in our system. Fig. S5 compares autophagosomal maturation state in the axonal subregions.

## Figure Legends

Figure S1. **Dynein effectors associate with LC3+ vesicles in the axon. (A)** Micrograph and line scan demonstrating LC3+ vesicles can colocalize with all three candidates simultaneously. Yellow dashed line indicates where line scan intensity measurement was taken; scale bar, 1 µm. **(B)** Schematic illustrating Proximity ligation assay (PLA) method. Bottom right, representative negative control showing few puncta are formed in the absence of primary antibody. All PLA graphs show dashed gray line for negative control representing *n* = 3 negative control neurons expressing that Halo-tagged candidate. (C-H) Representative micrographs and quantifications of PLA puncta for endogenous LC3 with Halo-tagged JIP1 **(C-D),** HAP1 **(E-F)**, and JIP3 **(G-H)** along the axon (dotted gray line). Arrows indicate proximity ligation assay (PLA) puncta. Scale bar, 10 µm. Dashed gray line, negative control (missing primary antibody). *n* = 9-10 neurons; one-way ANOVA demonstrates none show statistical significance between regions. **(I)** Immunoblotting of autophagosomal isolation illustrating enrichment of LC3 and Htt but not Golgi marker GM130. **(J)** Quantification of lipidated autophagosome-associated LC3-II isoform compared to the cytosolic non-lipidated LC3-I isoform showing the autophagosomal fraction (A) is enriched for autophagic vacuoles. *n* = 3 preps; two-tailed paired t test (*p* = 0.0254). **(K)** Quantification of enrichment in the autophagosomal fraction, displayed as relative to brain lysate (input). *n* = 3 preps; one-way ANOVA for each protein; GM130 (*p* = 0.2128), Htt (*p* = 0.0198). (L-O) Representative micrographs and quantifications of PLA puncta for endogenous dynein intermediate chain (DIC) with Halo-tagged HAP1 **(L-M)**, and JIP3 **(N-O)** along the axon. Symbols all same as (C-H). Bars throughout figure show mean ± SEM.

Figure S2. **Htt regulates autophagosomal motility in the mid-axon. (A)** Immunoblotting of PC12 cell lysates demonstrate KD efficiency of HAP1 and Htt siRNAs. Normalization factor (NF) determined using Revert^TM^ Total Protein Stain. **(B)** Immunoblot quantification for *n* = 4-5 repeats. **(C)** Representative kymographs from the mid-axon of a mock-transfected (control) neuron and a neuron transfected with Htt siRNA. (D-F) Quantification of LC3+ puncta motile behavior in the distal **(D)**, mid-**(E)**, and proximal **(F)** axonal subregions. *n* = 6-21 neurons. Two-way ANOVA for each with Sidak’s multiple comparisons test (Mid-axon Retrograde Mock v. Htt KD, *p* = 0.0076; Mid-axon Stationary/Bidirectional Mock v. Htt KD, *p* = 0.0132). (G-H) Representative kymograph **(G)** and quantification **(H)** from the mid-axon of a neuron transfected with Htt siRNA and siRNA-resistant mCh-Htt. Two-way ANOVA with Tukey’s multiple comparisons test (Retrograde Mock v. Htt KD, *p* = 0.0005=4; Stationary/Bidirectional Mock v. Htt KD, *p* = 0.0009). Bars throughout figure show mean ± SEM. Symbols (ns, not significant; *, *p* < 0.05; **, *p* < 0.01; ***, *p* < 0.001) indicate Sidak’s or Tukey’s test comparing to Mock. **Conservation of the *Glued* binding motif. (I)** Sequence alignment of HAP1 (human aa 280-445) across species. (J-K) Alignment of human HAP1 (aa 307-370) with known and putative dynein activating adaptors (human canonical sequences). The region is well-conserved with the TRAK and HOOK families **(J)** but less so with the BICD, SPINDLY, and EF-hand-containing (CRACR2a) families of dynein effectors **(K).** All alignments run with T-Coffee (ver. 11.00.d625267) default settings. Prepared for visualization by BoxShade (ExPAy, Swiss Institute of Bioinformatics) with RTF_new output. Black boxes, aa identical in ≥ 60% of sequences. Gray boxes, aa similar in ≥ 60% of sequences.

Figure S3. **HAP1 does not activate kinesin *in vitro*. (A-B)** Peroxisome recruitment assay showing peroxisome (PEX-GFP-DHFR) localization across a 20 min video. Addition of dimerizer tethers peroxisomes to HaloTag causing them to relocalize to the cell center **(A)** while addition of vehicle alone (DMSO) does not induce relocalization **(B)**. Left, initial (0 min) and concluding (25 min) stills of peroxisomes. Right, max time projection pseudocolored by frame (scale above). Yellow line indicates cell outline. Scale bar, 10 µm. **(C-D)** Quantification (C) and example micrographs (D) from kinesin MT recruitment TIRF assay wherein cell extracts expressing constitutively active N-terminal kinesin construct K560 or full-length kinesin-1 heavy chain (KHC) were added to Taxol-stabilized MT in a TIRF chamber with nonhydrolyzable ATP homolog AMPPNP. Intensity of TMR-labeled kinesin is measured across a 3 min video. Positive control K560 binds the MT robustly, while full-length KHC is autoinhibited and does not bind. Co-expression of HAP1 or HAP1 and Htt does not activate KHC, as seen by the lack of MT binding. *n* = 20 MT from 4 independent trials; one-way ANOVA (*p* < 0.0001) with Tukey’s multiple comparisons test (KHC v. K560, *p* = 0.0019; KHC v. KHC + HAP1, *p* = 0.9992; KHC v. KHC + HAP1 + Htt, *p* = 0.9833). Scale bar, 5 µm. **(E-F)** Blotting and quantification of cell extract MT pelleting assay for dynactin p150^Glued^ (p150; E) and dynein light intermediate chain 1 (LIC1; F) normalized to 5 µM MT only condition. *n* = 3-4 independent assays; one-way ANOVA (p150, *p* < 0.0001; DLIC, *p* = 0.0001) followed by Tukey’s multiple comparisons test. p150 is shown to bind MT in an ATP-dependent fashion (*p* = 0.0436). Increasing ionic strength (150 mM KCl) also decreased pellet fraction significantly without ATP (*p* = 0.0025) but with ATP increasing ionic strength did not change MT binding (*p* = 0.6624). DLIC is shown to bind MT in an ATP-dependent fashion (*p* = 0.0264). Increasing ionic strength (150 mM KCl) also decreased pellet fraction significantly without ATP (*p* = 0.0025) but with ATP increasing ionic strength did not change MT binding (*p* > 0.9999). HAP1 dynein-dynactin binding sites are not required for autophagosomal motility in the proximal axon. (G) Example kymograph illustrates the typical motility of EGFP-LC3+ puncta in the proximal axon in the presence of overexpressed Halo-HAP1^WT^. (H-I) Example kymographs (H) and quantification (I) showing no effect of HAP1 CC1 box mutants (HAP1^AAVV^, HAP1^ID^) on LC3+ puncta motility in the proximal axon. *n* = 10-13 neurons; two-way ANOVA with Bonferroni’s multiple comparisons test (Retrograde WT v. AAVV, *p* = 0.1337; Stationary/Bidirectional WT v. AAVV, *p* = 0.0900; Retrograde WT v. ID, *p* > 0.9999; Stationary/Bidirectional WT v. ID, *p* > 0.9999). (J-K) Example kymograph (J) and quantification (K) showing lack of effect of HAP1 *Glued* motif mutant (HAP1^EEAA^) on LC3+ puncta motility in the proximal axon. *n* = 9-10 neurons; two-way ANOVA with Bonferroni’s multiple comparisons test (Retrograde WT v. EEAA, *p* = 0.5408; Stationary/Bidirectional WT v. EEAA, *p* = 0.4316). (L-M) Example kymograph (L) and quantification (M) showing the HAP1 Spindly mutant (HAP1^TA^) does not have a dominant negative effect on LC3+ puncta motile behavior in the mid-axon. *n* = 16 neurons; two-way ANOVA with Bonferroni’s multiple comparisons test (Retrograde WT v. TA, *p* = 0.4635; Stationary/Bidirectional WT v. TA, *p* = 0.3500). **(N-O)** Example kymograph (N) and quantification (O) showing normal motile behavior in the proximal axon of cells transfected with both HAP1 siRNA and the HAP spindly mutant (HAP1^TA^). *n* = 10-12 neurons; two-way ANOVA with Bonferroni’s multiple comparisons test (Retrograde WT v. TA, *p* > 0.9999; Stationary/Bidirectional WT v. TA, *p* > 0.9999). Bars throughout figure show mean ± SEM. Symbols (ns, not significant; *, *p* < 0.05; **, *p* < 0.01; ***, *p* < 0.001) indicate Sidak’s or Tukey’s test comparing to Mock unless otherwise indicated.

Figure S4. **RILP is not a major regulator of autophagosomal motility in hippocampal axons. (A)** Time series and kymograph from separate LC3+ autophagosomes demonstrating comigration with RILP. Scale bars, 2 µm. **(B)** Quantification of LC3+ puncta comigrating with RILP in different subaxonal regions. *n* = 8-10 neurons; one-way ANOVA (*p* = 0.9577). (C-D) Example immunoblot **(C)** and quantification **(D)** of autophagosomal isolation illustrating enrichment of RILP on the autophagosomal surface. *n* = 3 preps; one-way ANOVA (*p* = 0.0039). (E-H) Representative micrographs and quantifications of PLA puncta for Halo-tagged RILP and endogenous LC3 **(E-F)** or endogenous dynein intermediate chain (DIC; **G-H)** along the axon (dotted gray line). Arrows indicate proximity ligation assay (PLA) puncta. Scale bar, 10 µm. Dashed gray line, negative control (missing primary antibody). *n* = 6-8 neurons; one-way ANOVA indicates neither shows statistically significant difference between regions. **(I)** Immunoblotting and quantification of PC12 cell lysates demonstrate KD efficiency of RILP siRNA. *n* = 4 repeats. Normalization factor (NF) determined using Revert^TM^ Total Protein Stain. **(J)** Representative kymographs from the proximal axon of a mock-transfected cell, a cell transfected with RILP siRNA, and a cell co-transfected with RILP siRNA and siRNA-resistant RILP-Halo. **(K)** Quantification of LC3+ puncta motile behavior in the distal, mid-, and proximal axonal subregions. *n* = 10-13 neurons. Two-way ANOVA for each with Bonferroni’s multiple comparisons test (Proximal Retrograde Mock v. RILP KD, *p* = 0.2416; Proximal Stationary/Bidirectional Mock v. RILP KD, *p* = 0.0237; Proximal Retrograde Mock v. Rescue, *p* = 0.0222; Proximal Stationary/Bidirectional Mock v. Rescue, *p* > 0.9999; Proximal Retrograde RILP KD v. Rescue, *p* > 0.9999; Proximal Stationary/Bidirectional RILP KD v. Rescue, *p* = 0.0141;). Colors represent retrograde-moving (cyan), stationary/bidirectional (magenta), and anterograde-moving (gray) puncta. Symbols indicate Bonferroni’s test comparing to Mock. Bars throughout figure show mean ± SEM. ns, not significant; *, *p* < 0.05; **, *p* < 0.01.

Figure S5. **Autophagosomes in central nervous system axons mature more slowly than those in peripheral nervous system axons.** (A-D) Example kymographs and quantification of comigrating Rab7 **(A-B)** or LAMP1 **(C-D)** with LC3 in different subaxonal regions. *n* = 10-12 neurons. Two-way ANOVA with Tukey’s multiple comparisons test (Rab7: ANOVA, *p* = 0.8648. LAMP1: ANOVA, *p* = 0.0343; Distal v. Mid-axon, *p* = 0.4166; Distal v. Proximal, *p* = 0.0264; Mid-axon v. Proximal, *p* = 0.3205). (E-F) Example kymographs **(E)** and quantification **(F)** of LC3+ puncta comigrating with LysoTracker^TM^ Deep Red dye in different subaxonal regions. LysoTracker specifically labels acidic organelles. *n* = 8-10 neurons. Two-way ANOVA with Tukey’s multiple comparisons test (Distal v. Mid-axon, *p* = 0.0709; Distal v. Proximal, *p* = 0.0280; Mid-axon v. Proximal, *p* = 0.8623). (G-H) Example micrographs with line scans **(G)** and quantification **(H)** demonstrating RILP association with autophagosomes of differing maturity. *n* = 8-9 neurons; some videos had exclusively mCh+EGFP or mCh only so the number of points on each graph may vary. Two-way ANOVA with Sidak’s multiple comparisons test (Mid-axon mCh+EGFP v. mCh only, *p* = 0.4894; Proximal mCh+EGFP v. mCh only, *p* = 0.0403). Dashed yellow lines indicate location of line scan.Scale bar, 1 µm. Bars throughout figure show mean ± SEM. ns, not significant; *, *p* < 0.05.

## References

1. Alam, M.S. 2018. Proximity Ligation Assay. Current Protocols in Immunology.

2. Arimoto, M., S.P. Koushika, B.C. Choudhary, C. Li, K. Matsumoto, and N. Hisamoto. 2011. The Caenorhabditis elegans JIP3 Protein UNC-16 Functions As an Adaptor to Link Kinesin-1 with Cytoplasmic Dynein. J. Neurosci. 31:2216–2224. doi:10.1523/JNEUROSCI.2653-10.2011.

3. Ayloo, S., J.E. Lazarus, A. Dodda, M. Tokito, E.M. Ostap, and E.L.F. Holzbaur. 2014. Dynactin functions as both a dynamic tether and brake during dynein-driven motility. Nat Commun. 5:4807. doi:10.1038/ncomms5807.

4. Ballister, E.R., S. Ayloo, D.M. Chenoweth, M.A. Lampson, and E.L.F. Holzbaur. 2015. Optogenetic control of organelle transport using a photocaged chemical inducer of dimerization. Curr Biol. 25:R407–R408. doi:10.1016/j.cub.2015.03.056.

5. Beeler, N., B.M. Riederer, G. Waeber, and A. Abderrahmani. 2009. Role of the JNK-interacting protein 1/islet brain 1 in cell degeneration in Alzheimer disease and diabetes. Brain Research Bulletin. 80:274–281. doi:10.1016/j.brainresbull.2009.07.006.

6. Brown, H.M., H.A.V. Epps, A. Goncharov, B.D. Grant, and Y. Jin. 2009. The JIP3 scaffold protein UNC-16 regulates RAB-5 dependent membrane trafficking at C. elegans synapses. Developmental Neurobiology. 69:174–190. doi:10.1002/dneu.20690.

7. Cai, Q., L. Lu, J.-H. Tian, Y.-B. Zhu, H. Qiao, and Z.-H. Sheng. 2010. Snapin-Regulated Late Endosomal Transport Is Critical for Efficient Autophagy-Lysosomal Function in Neurons. Neuron. 68:73–86. doi:10.1016/j.neuron.2010.09.022.

8. Cavalli, V., P. Kujala, J. Klumperman, and L.S.B. Goldstein. 2005. Sunday Driver links axonal transport to damage signaling. J. Cell Biol. 168:775–787. doi:10.1083/jcb.200410136.

9. Caviston, J.P., J.L. Ross, S.M. Antony, M. Tokito, and E.L. Holzbaur. 2007. Huntingtin facilitates dynein/dynactin-mediated vesicle transport. Proceedings of the National Academy of Sciences. 104:10045–10050.

10. Celestino, R., M.A. Henen, J.B. Gama, C. Carvalho, M. McCabe, D.J. Barbosa, A. Born, P.J. Nichols, A.X. Carvalho, R. Gassmann, and B. Vögeli. 2019. A transient helix in the disordered region of dynein light intermediate chain links the motor to structurally diverse adaptors for cargo transport. PLoS Biol. 17. doi:10.1371/journal.pbio.3000100.

11. Cheng, X.-T., B. Zhou, M.-Y. Lin, Q. Cai, and Z.-H. Sheng. 2015. Axonal autophagosomes recruit dynein for retrograde transport through fusion with late endosomes. J Cell Biol. 209:377–386. doi:10.1083/jcb.201412046.

12. Choudhary, B., M. Kamak, N. Ratnakaran, J. Kumar, A. Awasthi, C. Li, K. Nguyen, K. Matsumoto, N. Hisamoto, and S.P. Koushika. 2017. UNC-16/JIP3 regulates early events in synaptic vesicle protein trafficking via LRK-1/LRRK2 and AP complexes. PLOS Genetics. 13:e1007100. doi:10.1371/journal.pgen.1007100.

13. Cianfrocco, M.A., M.E. DeSantis, A.E. Leschziner, and S.L. Reck-Peterson. 2015. Mechanism and Regulation of Cytoplasmic Dynein. Annu. Rev. Cell Dev. Biol. 31:83–108. doi:10.1146/annurev-cellbio-100814-125438.

14. Cockburn, J.J.B., S.J. Hesketh, P. Mulhair, M. Thomsen, M.J. O’Connell, and M. Way. 2018. Insights into Kinesin-1 Activation from the Crystal Structure of KLC2 Bound to JIP3. Structure. 26:1486–1498.e6. doi:10.1016/j.str.2018.07.011.

15. Drerup, C.M., and A.V. Nechiporuk. 2013. JNK-Interacting Protein 3 Mediates the Retrograde Transport of Activated c-Jun N-Terminal Kinase and Lysosomes. PLOS Genetics. 9:e1003303. doi:10.1371/journal.pgen.1003303.

16. Elshenawy, M.M., J.T. Canty, L. Oster, L.S. Ferro, Z. Zhou, S.C. Blanchard, and A. Yildiz. 2019. Cargo adaptors regulate stepping and force generation of mammalian dynein–dynactin. Nature Chemical Biology. 15:1093–1101. doi:10.1038/s41589-019-0352-0.

17. Elshenawy, M.M., E. Kusakci, S. Volz, J. Baumbach, S.L. Bullock, and A. Yildiz. 2020. Lis1 activates dynein motility by modulating its pairing with dynactin. Nature Cell Biology. 22:570–578. doi:10.1038/s41556-020-0501-4.

18. Engelender, S., A.H. Sharp, V. Colomer, M.K. Tokito, A. Lanahan, P. Worley, E.L. Holzbaur, and C.A. Ross. 1997. Huntingtin-associated Protein 1 (HAP1) Interacts with the p150 Glued Bubunit of Dynactin. Human molecular genetics. 6:2205–2212.

19. Feng, Q., A.M. Gicking, and W.O. Hancock. 2020. Dynactin p150 promotes processive motility of DDB complexes by minimizing diffusional behavior of dynein. MBoC. 31:782–792. doi:10.1091/mbc.E19-09-0495.

20. Fu, M., and E.L.F. Holzbaur. 2013. JIP1 regulates the directionality of APP axonal transport by coordinating kinesin and dynein motors. J Cell Biol. 202:495–508. doi:10.1083/jcb.201302078.

21. Fu, M., and E.L.F. Holzbaur. 2014. Integrated regulation of motor-driven organelle transport by scaffolding proteins. Trends in Cell Biology. 24:564–574. doi:10.1016/j.tcb.2014.05.002.

22. Fu, M., J.J. Nirschl, and E.L.F. Holzbaur. 2014. LC3 Binding to the Scaffolding Protein JIP1 Regulates Processive Dynein-Driven Transport of Autophagosomes. Dev Cell. 29:577–590. doi:10.1016/j.devcel.2014.04.015.

23. Gama, J.B., C. Pereira, P.A. Simões, R. Celestino, R.M. Reis, D.J. Barbosa, H.R. Pires, C. Carvalho, J. Amorim, A.X. Carvalho, D.K. Cheerambathur, and R. Gassmann. 2017. Molecular mechanism of dynein recruitment to kinetochores by the Rod– Zw10–Zwilch complex and Spindly. J Cell Biol. 216:943–960. doi:10.1083/jcb.201610108.

24. Gassmann, R., A.J. Holland, D. Varma, X. Wan, F. Çivril, D.W. Cleveland, K. Oegema, E.D. Salmon, and A. Desai. 2010. Removal of Spindly from microtubule-attached kinetochores controls spindle checkpoint silencing in human cells. Genes Dev. 24:957–971. doi:10.1101/gad.1886810.

25. George, A.A., S. Hayden, G.R. Stanton, and S.E. Brockerhoff. 2016. Arf6 and the 5’phosphatase of synaptojanin 1 regulate autophagy in cone photoreceptors. BioEssays. 38:S119–S135. doi:10.1002/bies.201670913.

26. Gill, S.R., T.A. Schroer, I. Szilak, E.R. Steuer, M.P. Sheetz, and D.W. Cleveland. 1991. Dynactin, a conserved, ubiquitously expressed component of an activator of vesicle motility mediated by cytoplasmic dynein. J Cell Biol. 115:1639–1650.

27. Gowrishankar, S., Y. Wu, and S.M. Ferguson. 2017. Impaired JIP3-dependent axonal lysosome transport promotes amyloid plaque pathology. J Cell Biol. 216:3291–3305. doi:10.1083/jcb.201612148.

28. Grotjahn, D.A., S. Chowdhury, Y. Xu, R.J. McKenney, T.A. Schroer, and G.C. Lander. 2018. Cryo-electron tomography reveals that dynactin recruits a team of dyneins for processive motility. Nat Struct Mol Biol. 25:203–207. doi:10.1038/s41594-018-0027-7.

29. Gunawardena, S., L.-S. Her, R.G. Brusch, R.A. Laymon, I.R. Niesman, B. Gordesky-Gold, L. Sintasath, N.M. Bonini, and L.S.B. Goldstein. 2003. Disruption of Axonal Transport by Loss of Huntingtin or Expression of Pathogenic PolyQ Proteins in Drosophila. Neuron. 40:25–40. doi:10.1016/S0896-6273(03)00594-4.

30. Hancock, W.O. 2014. Bidirectional cargo transport: moving beyond tug of war. Nat. Rev. Mol. Cell Biol. 15:615–628. doi:10.1038/nrm3853.

31. Hara, T., K. Nakamura, M. Matsui, A. Yamamoto, Y. Nakahara, R. Suzuki-Migishima, M. Yokoyama, K. Mishima, I. Saito, H. Okano, and N. Mizushima. 2006. Suppression of basal autophagy in neural cells causes neurodegenerative disease in mice. Nature. 441:885–889. doi:10.1038/nature04724.

32. Heidemann, S.R., J.M. Landers, and M.A. Hamborg. 1981. Polarity orientation of axonal microtubules. J Cell Biol. 91:661–665. doi:10.1083/jcb.91.3.661.

33. Helbecque, N., A. Abderrhamani, L. Meylan, B. Riederer, V. Mooser, J. Miklossy, J. Delplanque, P. Boutin, P. Nicod, J.-A. Haefliger, D. Cottel, P. Amouyel, P. Froguel, and G. Waeber. 2003. Islet-brain1/C-Jun N-terminal kinase interacting protein-1 (IB1/JIP-1) promoter variant is associated with Alzheimer’s disease. Molecular Psychiatry. 8:413–422. doi:10.1038/sj.mp.4001344.

34. Hendricks, A.G., E. Perlson, J.L. Ross, H.W. Schroeder, M. Tokito, and E.L.F. Holzbaur. 2010. Motor Coordination via a Tug-of-War Mechanism Drives Bidirectional Vesicle Transport. Current Biology. 20:697–702. doi:10.1016/j.cub.2010.02.058.

35. Henrichs, V., L. Grycova, C. Barinka, Z. Nahacka, J. Neuzil, S. Diez, J. Rohlena, M. Braun, and Z. Lansky. 2020. Mitochondria-adaptor TRAK1 promotes kinesin-1 driven transport in crowded environments. Nature Communications. 11:3123. doi:10.1038/s41467-020-16972-5.

36. Hill, S.E., K.J. Kauffman, M. Krout, J.E. Richmond, T.J. Melia, and D.A. Colón-Ramos. 2019. Maturation and Clearance of Autophagosomes in Neurons Depends on a Specific Cysteine Protease Isoform, ATG-4.2. Developmental Cell. 49:251–266.e8. doi:10.1016/j.devcel.2019.02.013.

37. Hodgkinson, J.L., C. Peters, S.A. Kuznetsov, and W. Steffen. 2005. Three-dimensional reconstruction of the dynactin complex by single-particle image analysis. PNAS. 102:3667–3672. doi:10.1073/pnas.0409506102.

38. Itakura, E., C. Kishi-Itakura, and N. Mizushima. 2012. The Hairpin-type Tail-Anchored SNARE Syntaxin 17 Targets to Autophagosomes for Fusion with Endosomes/Lysosomes. Cell. 151:1256–1269. doi:10.1016/j.cell.2012.11.001.

39. Khobrekar, N.V., S. Quintremil, T.J. Dantas, and R.B. Vallee. 2020. The Dynein Adaptor RILP Controls Neuronal Autophagosome Biogenesis, Transport, and Clearance. Developmental Cell. doi:10.1016/j.devcel.2020.03.011.

40. Komatsu, M., S. Waguri, T. Chiba, S. Murata, J. Iwata, I. Tanida, T. Ueno, M. Koike, Y. Uchiyama, E. Kominami, and K. Tanaka. 2006. Loss of autophagy in the central nervous system causes neurodegeneration in mice. Nature. 441:880–884. doi:10.1038/nature04723.

41. Kulkarni, A., J. Chen, and S. Maday. 2018. Neuronal autophagy and intercellular regulation of homeostasis in the brain. Current Opinion in Neurobiology. 51:29–36. doi:10.1016/j.conb.2018.02.008.

42. Kumar, S., Y. Zhou, and M. Plamann. 2001. Dynactin–membrane interaction is regulated by the C-terminal domains of p150Glued. EMBO Rep. 2:939–944. doi:10.1093/embo-reports/kve202.

43. Kunwar, A., S.K. Tripathy, J. Xu, M.K. Mattson, P. Anand, R. Sigua, M. Vershinin, R.J. McKenney, C.C. Yu, A. Mogilner, and S.P. Gross. 2011. Mechanical stochastic tug-of-war models cannot explain bidirectional lipid-droplet transport. PNAS. 108:18960– 18965. doi:10.1073/pnas.1107841108.

44. Kural, C., H. Kim, S. Syed, G. Goshima, V.I. Gelfand, and P.R. Selvin. 2005. Kinesin and Dynein Move a Peroxisome in Vivo: A Tug-of-War or Coordinated Movement? Science. 308:1469–1472. doi:10.1126/science.1108408.

45. Lau, C.K., F.J. O’Reilly, B. Santhanam, S.E. Lacey, J. Rappsilber, and A.P. Carter. 2020. Cryo-EM reveals the complex architecture of dynactin’s shoulder and pointed end. bioRxiv. 2020.07.16.206359. doi:10.1101/2020.07.16.206359.

46. Lee, I.-G., S.E. Cason, S.S. Alqassim, E.L.F. Holzbaur, and R. Dominguez. 2020. A tunable LIC1-adaptor interaction modulates dynein activity in a cargo-specific manner. Nature Communications. Accepted.

47. Lee, I.-G., M.A. Olenick, M. Boczkowska, C. Franzini-Armstrong, E.L.F. Holzbaur, and R. Dominguez. 2018. A conserved interaction of the dynein light intermediate chain with dynein-dynactin effectors necessary for processivity. Nature Communications. 9:986. doi:10.1038/s41467-018-03412-8.

48. Li, S.-H., C.-A. Gutekunst, S.M. Hersch, and X.-J. Li. 1998. Interaction of huntingtin-associated protein with dynactin P150Glued. Journal of Neuroscience. 18:1261– 1269.

49. Li, X.J., A.H. Sharp, S.H. Li, T.M. Dawson, S.H. Snyder, and C.A. Ross. 1996. Huntingtin-associated protein (HAP1): discrete neuronal localizations in the brain resemble those of neuronal nitric oxide synthase. Proc Natl Acad Sci U S A. 93:4839–4844.

50. Ludwiczak, J., A. Winski, K. Szczepaniak, V. Alva, and S. Dunin-Horkawicz. 2019. DeepCoil—a fast and accurate prediction of coiled-coil domains in protein sequences. Bioinformatics. 35:2790–2795. doi:10.1093/bioinformatics/bty1062.

51. Lumsden, A.L., R.L. Young, N. Pezos, and D.J. Keating. 2016. Huntingtin-associated protein 1: Eutherian adaptation from a TRAK-like protein, conserved gene promoter elements, and localization in the human intestine. BMC Evol Biol. 16. doi:10.1186/s12862-016-0780-3.

52. Lupas, A., M.V. Dyke, and J. Stock. 1991. Predicting coiled coils from protein sequences. Science. 252:1162–1164. doi:10.1126/science.252.5009.1162.

53. Maday, S., K.E. Wallace, and E.L.F. Holzbaur. 2012. Autophagosomes initiate distally and mature during transport toward the cell soma in primary neurons. J Cell Biol. 196:407–417. doi:10.1083/jcb.201106120.

54. Marchesin, V., A. Castro-Castro, C. Lodillinsky, A. Castagnino, J. Cyrta, H. Bonsang-Kitzis, L. Fuhrmann, M. Irondelle, E. Infante, G. Montagnac, F. Reyal, A. Vincent-Salomon, and P. Chavrier. 2015. ARF6–JIP3/4 regulate endosomal tubules for MT1-MMP exocytosis in cancer invasion. J Cell Biol. 211:339–358. doi:10.1083/jcb.201506002.

55. Matanis, T., A. Akhmanova, P. Wulf, E. Del Nery, T. Weide, T. Stepanova, N. Galjart, F. Grosveld, B. Goud, C.I. De Zeeuw, A. Barnekow, and C.C. Hoogenraad. 2002. Bicaudal-D regulates COPI-independent Golgi-ER transport by recruiting the dynein-dynactin motor complex. Nat Cell Biol. 4:986–992. doi:10.1038/ncb891.

56. Mauvezin, C., P. Nagy, G. Juhász, and T.P. Neufeld. 2015. Autophagosome–lysosome fusion is independent of V-ATPase-mediated acidification. Nat Commun. 6:1–14. doi:10.1038/ncomms8007.

57. McGuire, J.R., J. Rong, S.-H. Li, and X.-J. Li. 2006. Interaction of Huntingtin-associated Protein-1 with Kinesin Light Chain. doi:10.1074/jbc.M509806200.

58. McKenney, R.J., W. Huynh, M.E. Tanenbaum, G. Bhabha, and R.D. Vale. 2014. Activation of cytoplasmic dynein motility by dynactin-cargo adapter complexes. Science. 345:337–341. doi:10.1126/science.1254198.

59. Montagnac, G., J.-B. Sibarita, S. Loubéry, L. Daviet, M. Romao, G. Raposo, and P. Chavrier. 2009. ARF6 Interacts with JIP4 to Control a Motor Switch Mechanism Regulating Endosome Traffic in Cytokinesis. Current Biology. 19:184–195. doi:10.1016/j.cub.2008.12.043.

60. Morfini, G.A., Y.-M. You, S.L. Pollema, A. Kaminska, K. Liu, K. Yoshioka, B. Björkblom, E.T. Coffey, C. Bagnato, D. Han, C.-F. Huang, G. Banker, G. Pigino, and S.T. Brady. 2009. Pathogenic Huntingtin Inhibits Fast Axonal Transport by Activating JNK3 and Phosphorylating Kinesin. Nat Neurosci. 12:864–871. doi:10.1038/nn.2346.

61. Müller, M.J.I., S. Klumpp, and R. Lipowsky. 2008. Tug-of-war as a cooperative mechanism for bidirectional cargo transport by molecular motors. PNAS. 105:4609– 4614. doi:10.1073/pnas.0706825105.

62. Neisch, A.L., T.P. Neufeld, and T.S. Hays. 2017. A STRIPAK complex mediates axonal transport of autophagosomes and dense core vesicles through PP2A regulation. J Cell Biol. 216:441–461. doi:10.1083/jcb.201606082.

63. Ochaba, J., T. Lukacsovich, G. Csikos, S. Zheng, J. Margulis, L. Salazar, K. Mao, A.L. Lau, S.Y. Yeung, S. Humbert, F. Saudou, D.J. Klionsky, S. Finkbeiner, S.O. Zeitlin, J.L. Marsh, D.E. Housman, L.M. Thompson, and J.S. Steffan. 2014. Potential function for the Huntingtin protein as a scaffold for selective autophagy. Proceedings of the National Academy of Sciences. 111:16889–16894. doi:10.1073/pnas.1420103111.

64. Olenick, M.A., R. Dominguez, and E.L.F. Holzbaur. 2019. Dynein activator Hook1 is required for trafficking of BDNF-signaling endosomes in neurons. J Cell Biol. 218:220–233. doi:10.1083/jcb.201805016.

65. Olenick, M.A., and E.L.F. Holzbaur. 2019. Dynein activators and adaptors at a glance. J Cell Sci. 132. doi:10.1242/jcs.227132.

66. Olenick, M.A., M. Tokito, M. Boczkowska, R. Dominguez, and E.L.F. Holzbaur. 2016. Hook Adaptors Induce Unidirectional Processive Motility by Enhancing the Dynein-Dynactin Interaction. J. Biol. Chem. 291:18239–18251. doi:10.1074/jbc.M116.738211.

67. Qiu, R., J. Zhang, and X. Xiang. 2018. p25 of the dynactin complex plays a dual role in cargo binding and dynactin regulation. J. Biol. Chem. 293:15606–15619. doi:10.1074/jbc.RA118.004000.

68. Reck-Peterson, S.L., W.B. Redwine, R.D. Vale, and A.P. Carter. 2018. The cytoplasmic dynein transport machinery and its many cargoes. Nature Reviews Molecular Cell Biology. 1. doi:10.1038/s41580-018-0004-3.

69. Saudou, F., and S. Humbert. 2016. The Biology of Huntingtin. Neuron. 89:910–926. doi:10.1016/j.neuron.2016.02.003.

70. Schlager, M.A., A. Serra-Marques, I. Grigoriev, L.F. Gumy, M. Esteves da Silva, P.S. Wulf, A. Akhmanova, and C.C. Hoogenraad. 2014. Bicaudal D Family Adaptor Proteins Control the Velocity of Dynein-Based Movements. Cell Reports. 8:1248– 1256. doi:10.1016/j.celrep.2014.07.052.

71. Schroeder, C.M., and R.D. Vale. 2016. Assembly and activation of dynein–dynactin by the cargo adaptor protein Hook3. J Cell Biol. 214:309–318. doi:10.1083/jcb.201604002.

72. Smith, J.J., and J.D. Aitchison. 2013. Peroxisomes take shape. Nat Rev Mol Cell Biol. 14:803–817. doi:10.1038/nrm3700.

73. Stavoe, A.K.H., S.E. Hill, D.H. Hall, and D.A. Colón-Ramos. 2016. KIF1A/UNC-104 transports ATG-9 to regulate neurodevelopment and autophagy at synapses. Dev Cell. 38:171–185. doi:10.1016/j.devcel.2016.06.012.

74. Strømhaug, P.E., T.O. Berg, M. Fengsrud, and P.O. Seglen. 1998. Purification and characterization of autophagosomes from rat hepatocytes. Biochem J. 335:217–224.

75. Sun, F., C. Zhu, R. Dixit, and V. Cavalli. 2011. Sunday Driver/JIP3 binds kinesin heavy chain directly and enhances its motility. EMBO J. 30:3416–3429. doi:10.1038/emboj.2011.229.

76. Trigg, J., K. Gutwin, A.E. Keating, and B. Berger. 2011. Multicoil2: Predicting Coiled Coils and Their Oligomerization States from Sequence in the Twilight Zone. PLOS ONE. 6:e23519. doi:10.1371/journal.pone.0023519.

77. Truebestein, L., and T.A. Leonard. 2016. Coiled-coils: The long and short of it. BioEssays. 38:903–916. doi:10.1002/bies.201600062.

78. Twelvetrees, A.E., F. Lesept, E.L.F. Holzbaur, and J.T. Kittler. 2019. The adaptor proteins HAP1a and GRIP1 collaborate to activate the kinesin-1 isoform KIF5C. J Cell Sci. 132. doi:10.1242/jcs.215822.

79. Twelvetrees, A.E., E.Y. Yuen, I.L. Arancibia-Carcamo, A.F. MacAskill, P. Rostaing, M.J. Lumb, S. Humbert, A. Triller, F. Saudou, Z. Yan, and J.T. Kittler. 2010. Delivery of GABAARs to Synapses Is Mediated by HAP1-KIF5 and Disrupted by Mutant Huntingtin. Neuron. 65:53–65. doi:10.1016/j.neuron.2009.12.007.

80. Urnavicius, L., C.K. Lau, M.M. Elshenawy, E. Morales-Rios, C. Motz, A. Yildiz, and A.P. Carter. 2018. Cryo-EM shows how dynactin recruits two dyneins for faster movement. Nature. 554:202–206. doi:10.1038/nature25462.

81. Urnavicius, L., K. Zhang, A.G. Diamant, C. Motz, M.A. Schlager, M. Yu, N.A. Patel, C.V. Robinson, and A.P. Carter. 2015. The structure of the dynactin complex and its interaction with dynein. Science. 347:1441–1446. doi:10.1126/science.aaa4080.

82. Vilela, F., C. Velours, M. Chenon, M. Aumont-Nicaise, V. Campanacci, A. Thureau, O. Pylypenko, J. Andreani, P. Llinas, and J. Ménétrey. 2019. Structural characterization of the RH1-LZI tandem of JIP3/4 highlights RH1 domains as a cytoskeletal motor-binding motif. Sci Rep. 9:1–15. doi:10.1038/s41598-019-52537-3.

83. Vincent, T.L., P.J. Green, and D.N. Woolfson. 2013. LOGICOIL—multi-state prediction of coiled-coil oligomeric state. Bioinformatics. 29:69–76. doi:10.1093/bioinformatics/bts648.

84. Waeber, G., J. Delplanque, C. Bonny, V. Mooser, M. Steinmann, C. Widmann, A. Maillard, J. Miklossy, C. Dina, E.H. Hani, G. Waeber, J. Delplanque, N. Vionnet, P. Nicod, P. Boutin, and P. Froguel. 2000. The gene MAPK8IP1, encoding islet-brain-1, is a candidate for type 2 diabetes. Nature Genetics. 24:291–295. doi:10.1038/73523.

85. Weiss, K.R., and J.T. Littleton. 2016. Characterization of axonal transport defects in Drosophila Huntingtin mutants. J Neurogenet. 30:212–221. doi:10.1080/01677063.2016.1202950.

86. White, J.A., E. Anderson, K. Zimmerman, K.H. Zheng, R. Rouhani, and S. Gunawardena. 2015. Huntingtin differentially regulates the axonal transport of a sub-set of Rab-containing vesicles *in vivo*. Human Molecular Genetics. 24:7182–7195. doi:10.1093/hmg/ddv415.

87. Wong, Y.C., and E.L.F. Holzbaur. 2014. The Regulation of Autophagosome Dynamics by Huntingtin and HAP1 Is Disrupted by Expression of Mutant Huntingtin, Leading to Defective Cargo Degradation. J. Neurosci. 34:1293–1305. doi:10.1523/JNEUROSCI.1870-13.2014.

88. Wong, Y.C., and E.L.F. Holzbaur. 2015. Autophagosome dynamics in neurodegeneration at a glance. J. Cell. Sci. 128:1259–1267. doi:10.1242/jcs.161216.

89. Zhang, K., H.E. Foster, A. Rondelet, S.E. Lacey, N. Bahi-Buisson, A.W. Bird, and A.P. Carter. 2017. Cryo-EM Reveals How Human Cytoplasmic Dynein Is Auto-inhibited and Activated. Cell. 169:1303–1314.e18. doi:10.1016/j.cell.2017.05.025.

