## Supplementary figures and images for "Sequential dynein effectors regulate axonal autophagosome motility in a maturation-dependent pathway"

### Supplemental Figure 1

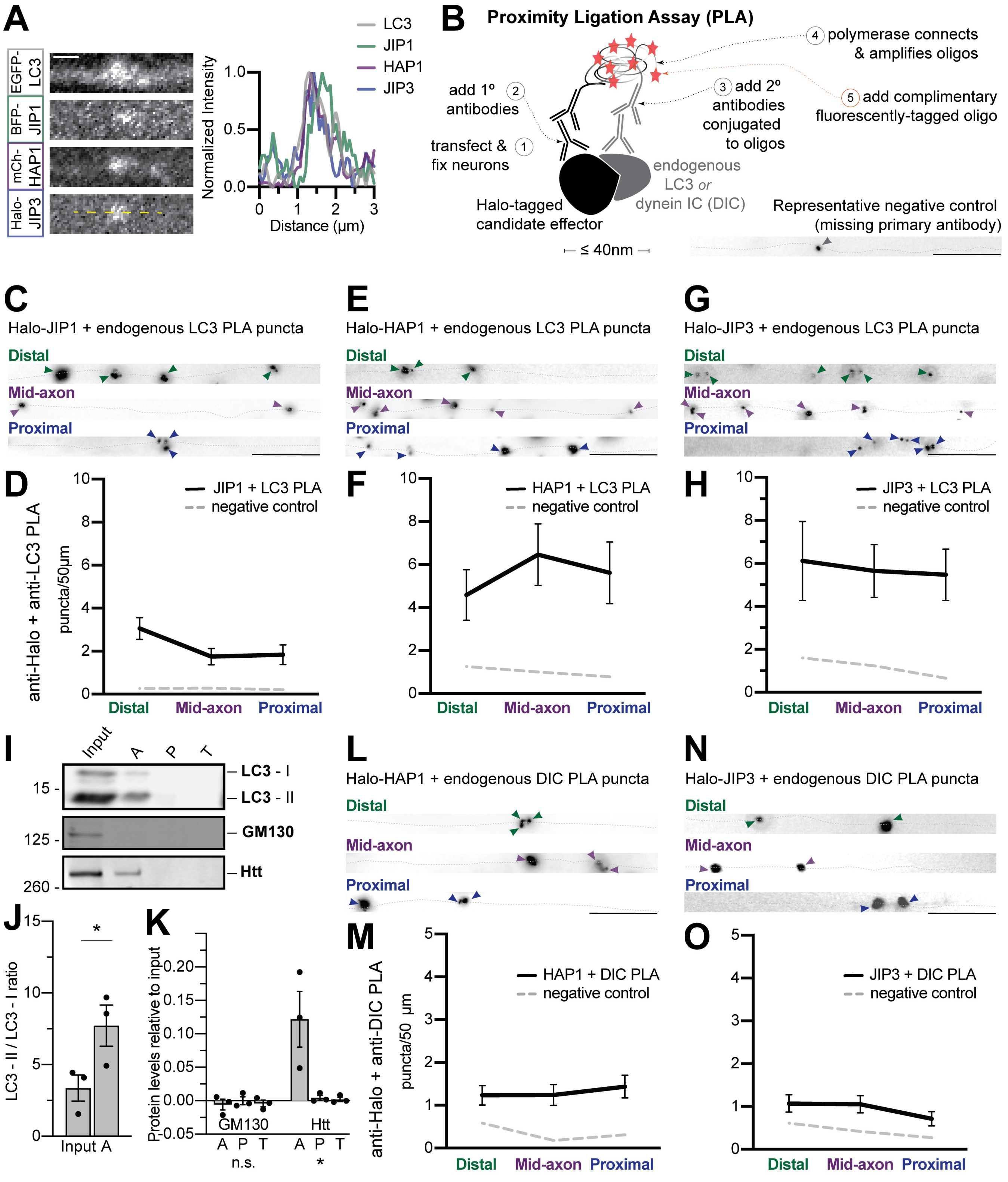

### Supplemental Figure 2

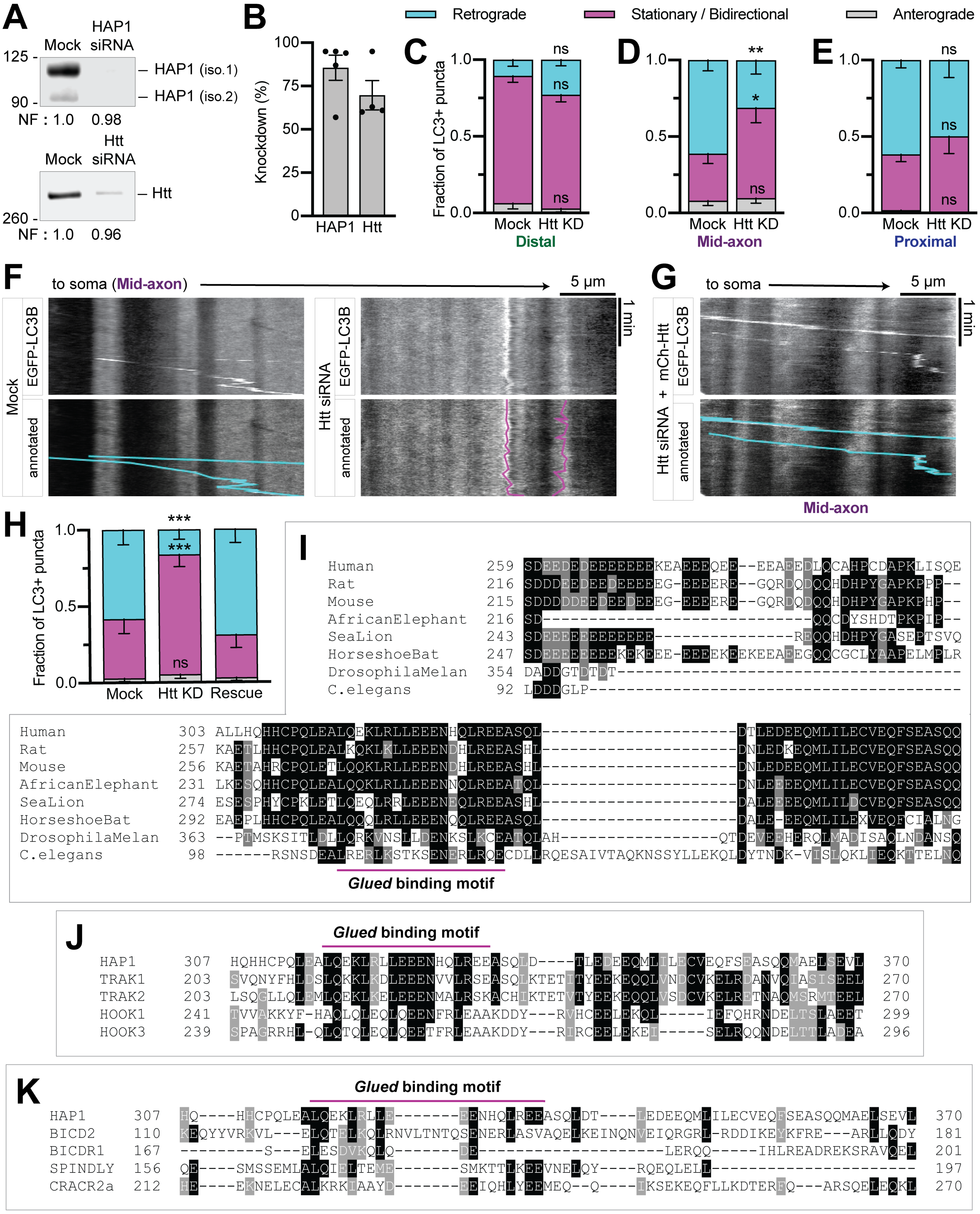

### Supplemental Figure 3

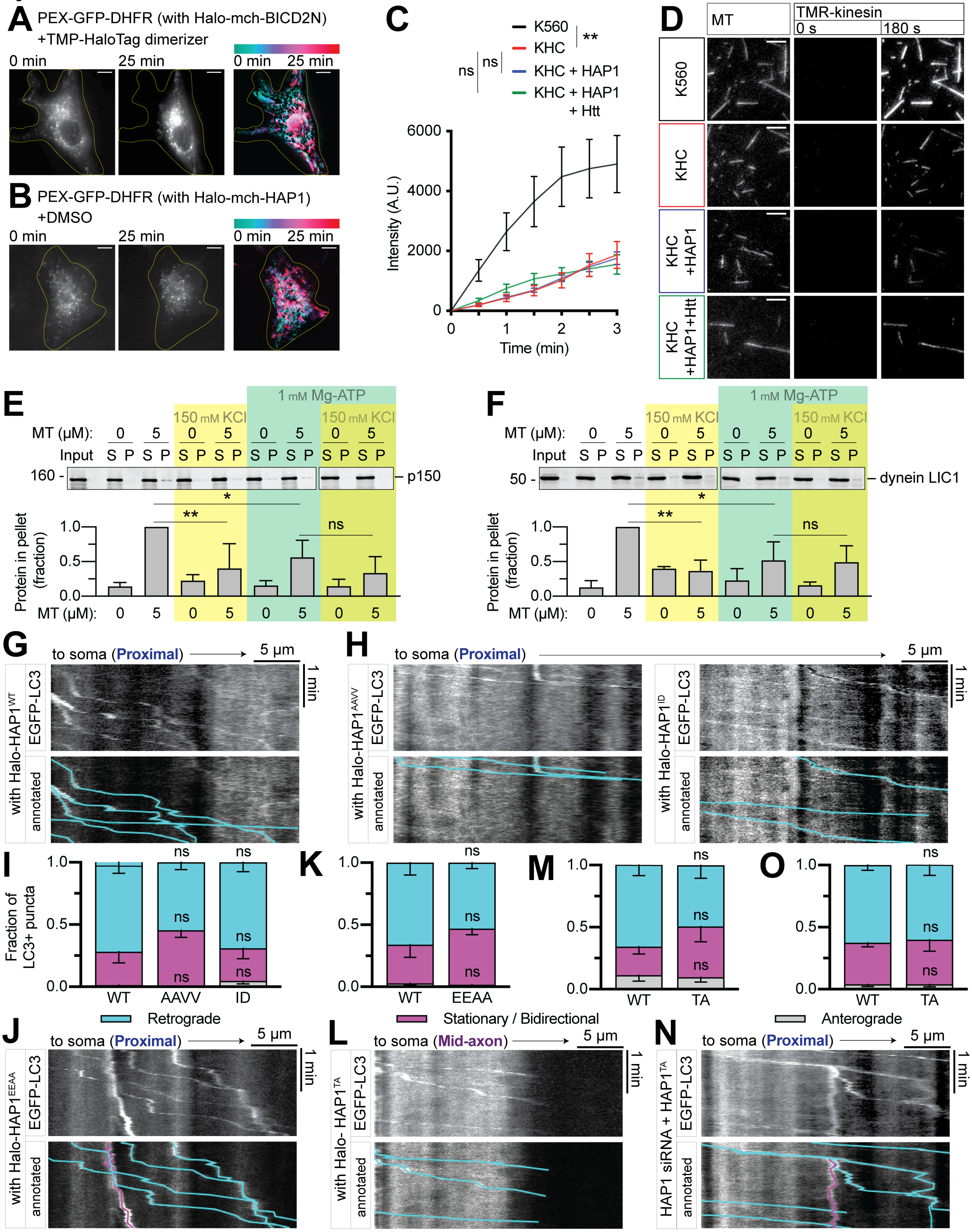

### Supplemental Figure 4

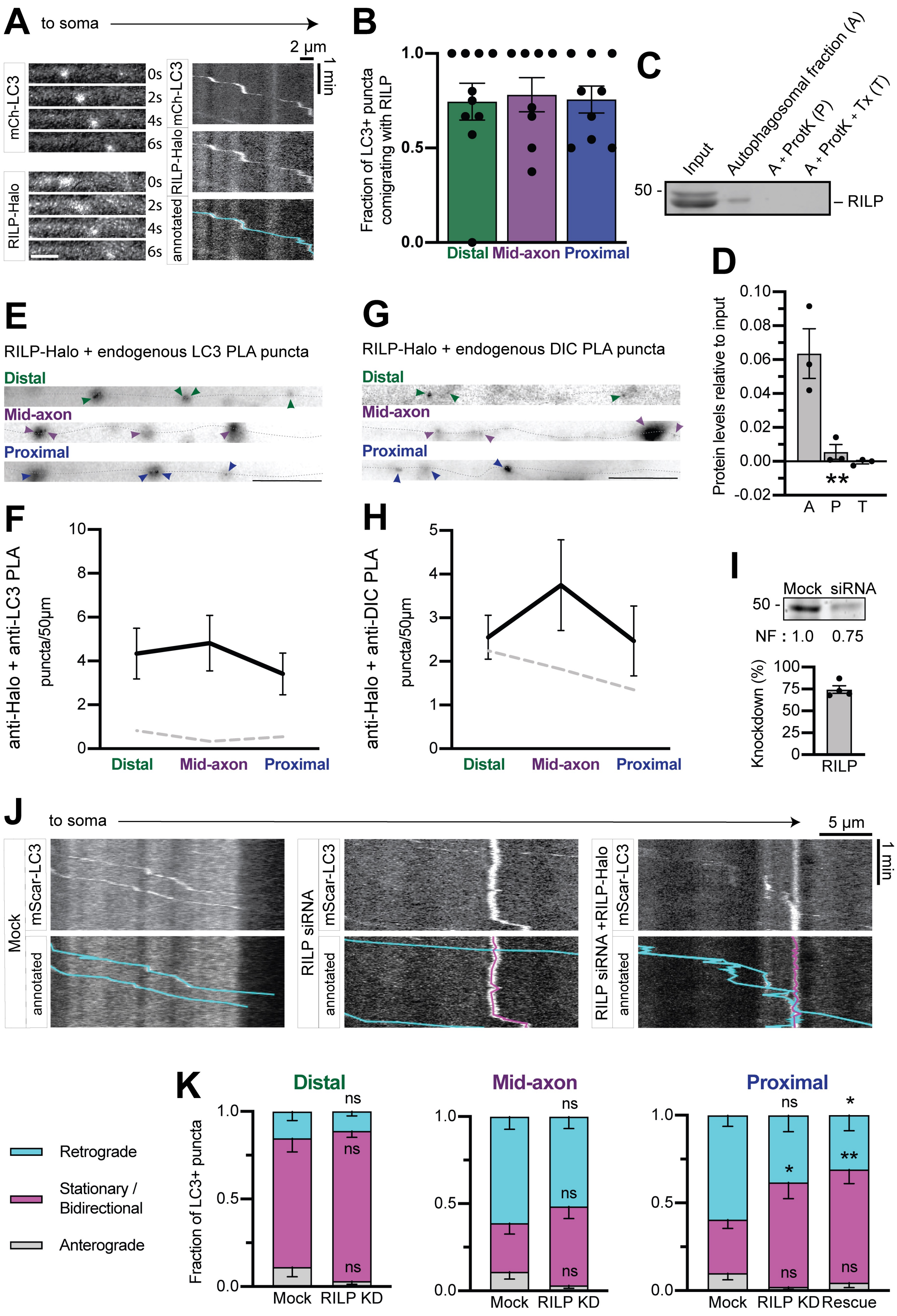

### Supplemental Figure 5

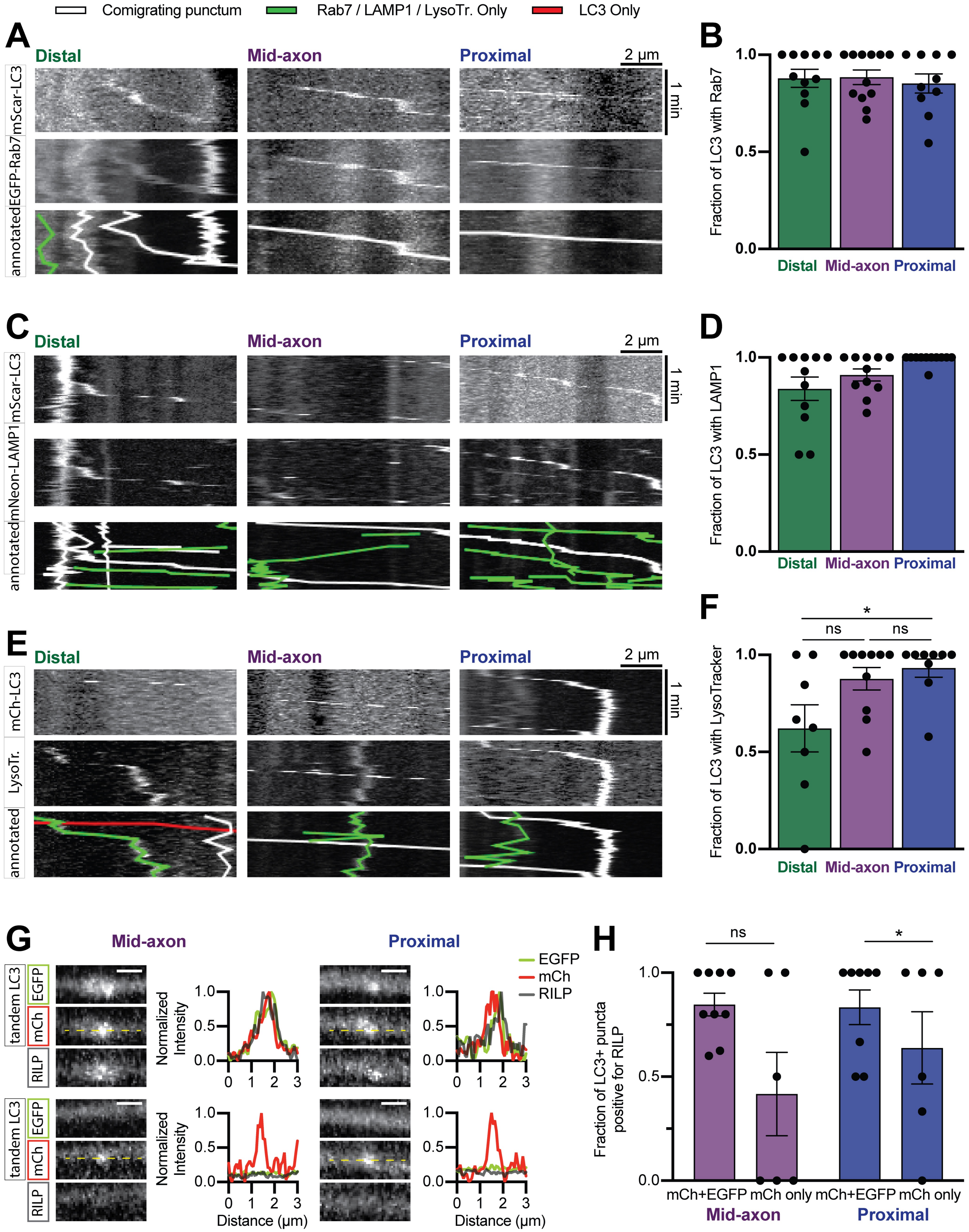
